# Paleogenomic time series reveal past population dynamics and their relevance for the conservation of the Alpine ibex

**DOI:** 10.64898/2026.09.28.755134

**Authors:** Xenia Wietlisbach, Mathieu Robin, Andreas Füglistaler, Giada Ferrari, Paul Cloarec-Pioffet, Andreas Zechner, Christophe Griggo, Michel Blant, Walter Imhof, Martin Trüssel, Jean-Marc Petillon, Gunther Gressman, Love Dalén, Daniel Wegmann, Christine Grossen

## Abstract

Endangered species often suffer from genetic threats such as inbreeding depression, the accumulation of mutation load and the general loss of genetic diversity, which may deprive them from future adaptation. Since the genetic make-up of a species is strongly affected by its evolutionary past, however, interspecific comparisons are difficult and absolute levels of genetic diversity may be a relatively poor predictor of extinction risk. To better interpret current levels of genetic diversity of actively managed species, the comparison of present-day genomic data to that of ancient material pre-dating recent anthropogenic population declines has been advocated, but remains rarely done at the evolutionary time scales necessary to understand past population dynamics. Here we demonstrate the usefulness of deep genomic time series through 40 high quality ancient genomes of the Alpine ibex, which faced near-extinction in the 19th century but has since been reintroduced across the Alps and recovered remarkably. The samples, which span the last 14,000 years, evidence strong and stable population structure at the continental scale, paired with frequent extinction and recolonization events at the local scale. These metapopulation dynamics resulted in small and largely isolated populations, with present-day, reintroduced populations only having 20% to 40% less neutral genetic diversity and 3% higher mutation load, likely due to efficient purging in the past. At the species level, however, the loss of genetic diversity was dramatic and reduced to the small fraction that segregated within the only surviving population in the Gran Paradiso area. Our analyses of ancient metapopulation dynamics in Alpine ibex underscore the importance of a healthy genetic pool for successful recolonization and long-term survival, as well as the real possibility of regional extinctions.

## Introduction

As many wild species, the charismatic Alpine ibex suffered from severe hunting pressure during the Early Modern Period, not least because essentially all its body parts were credited with healing properties. By the mid 19th century, a single population with less than 100 individuals had survived in the Gran Paradiso area, Italy, where the italian King Vittorio Emmanuelle II, a passionate hunter himself, established a royal hunting reserve and personally ensured their rigorous protection. In Switzerland, where the Alpine ibex was an essential part of alpine identity, the last Alpine ibex was shot at the beginning of the 19th century. As early as the mid 19th century, the Swiss government started serious efforts to procure individuals from the Italian king for reintroduction. Since the king consistently declined, the government later sponsored the hiring of poachers, who smuggled Alpine ibex kids to Switzerland from 1906 onwards. This resulted in one of the world’s first scientifically managed and exceptionally well documented reintroduction projects, during which the initial individuals were first raised and bred in captivity to create viable wild populations, from which additional populations continue to be stocked (Biebach & Keller, 2009; Grossen et al., 2018).

The genetic footprint of the past anthropogenic bottleneck is well visible among present-day Alpine ibex: they have very low levels of genome-wide diversity, comparable to that of the endangered Amur tiger, show high levels of inbreeding, in some cases consistent with matings among cousins, show an accumulation of deleterious mutation load (Grossen et al., 2020) and face hybridisation with, and introgression from, its domestic relative, the domestic goat (Brambilla et al., 2024; Grossen et al., 2014). Such footprints may prove fatal, particularly because inbreeding may lead to the expression of deleterious recessive alleles, as is well documented for the California condor (Robinson et al., 2021), the Eurasian lynx (Niehaus et al., 2025) or the Hawaiian crow (Kyriazis et al., 2026), among other recently bottlenecked species. While no severe genetic defects have been documented in Alpine ibex to date, higher inbreeding appears to slow population growth (Bozzuto et al., 2019) and lead to reduced horn growth (Brambilla et al., 2015). In addition, low diversity at immune-relevant genes was associated with increased disease susceptibility (Brambilla et al., 2018), and may explain the rapid introgression of domestic goat alleles at the MHC locus (Grossen et al., 2014). Nonetheless, the reintroduction of the Alpine ibex was remarkably successful: About 120 years or 15 generations after captive breeding started, the species is listed as Least Concern by the IUCN and more than 50’000 individuals roam in many populations spread across the entire European Alps (Brambilla et al., 2020). This shows that while reduced genetic diversity and population fragmentation is a common concern for threatened species (Willi et al., 2022), some species appear more resilient than others, at least in the short-term (Abson et al., 2026; Teixeira & Huber, 2021).

Given its well documented and well studied reintroduction history, as well as the availability of ample fossil material, the Alpine ibex presents an excellent opportunity to better understand how the deep genetic history may affect reintroduction success of threatened species. Alpine ibex depend on rocky areas and mountain meadows and only rarely traverse forests (Grignolio et al., 2003, 2007). This narrow niche between the upper timberline and mountain tops or glacier boundaries has continuously shifted upwards since the last glacial maximum, and likely only supported small and isolated populations for millenia. Here, we sequenced 40 ancient genomes of Alpine ibex spanning a time-series across the last 144,000 years to elucidate their deep demographic history, to understand how their ecological niche shaped their natural genetic makeup, and to investigate if that genetic legacy paved the way for their particularly fast recovery after the anthropogenically induced bottleneck.

## Results

We generated whole genome sequence data from teeth and bone material of 44 ancient Alpine ibex individuals from four key geographic regions of the current species distribution range (Figure 1A): i) 23 samples from the cave Tempiette in the Chartreuse region of France (Ch) (Griggo et al., 2019a, 2019b), ii) five samples from the the cave Giétroz in the Valais, Switzerland (Vs) (Reynaud Savioz et al., 2018; Reynaud Savioz & Blant, 2021), iii) 14 samples from eight caves and one glacier located in central Switzerland (CS), and iv) two samples from a cave and a glacier located in Austria (At). For comparison, we further generated whole genome sequence data of one ancient Iberian ibex from the Ossau valley in France. All sampled individuals were AMS-^14^C dated to between 19442 and 904 BP (Figure 1B, Supplementary Figure S1, Supplementary Table S1). We additionally sequenced 16 historic specimens from museums and collections, which were labeled as Alpine ibex and collected between 1695 and 1919 CE, based on available records. The sequence data of all ancient and historic samples showed signals of post-mortem damage, but varied greatly in endogenous DNA content from 13% to 74% (Supplementary Figure S2, Supplementary Table S1).

**Figure 1:**
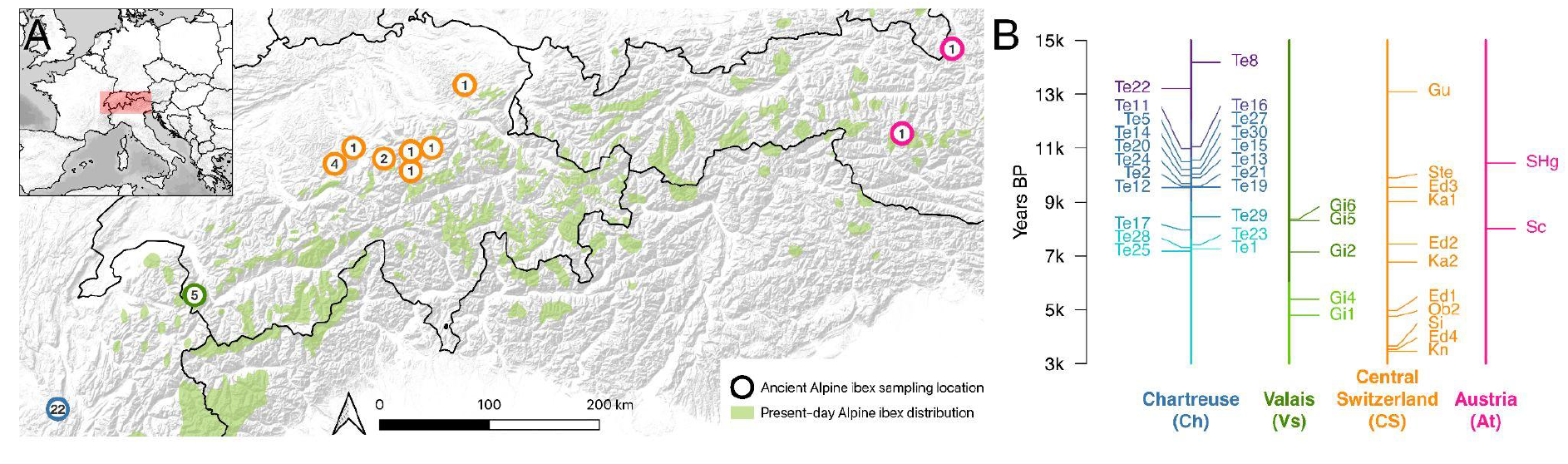
The geographic and temporal distribution of ancient Alpine ibex samples. A) Current geographic distribution of Alpine ibex (light green) and sampling locations with information on sample size (number) and geographic region (circle color): blue Chartreuse (Ch, France), green Valais (Vs, Switzerland), orange Central Switzerland (CS) and pink Austria (At). B) Temporal distribution (AMS-14C) of samples from each of the four geographic regions, colored as in (A). Two samples with very low depth (Str, Fi) and two samples that are likely duplicates (Ob2 and Te26) were removed and are not shown. See also Supplementary Table S1.

As indicated by extremely low genetic distance estimates (<10^-5^, 100-times lower than any other pair), very high kinship coefficients (KING > 0.48, Supplementary Figure S3) and virtually identical 14C dates (Supplementary Table S1), two pairs of ancient samples (Ob1 and Ob2; Te26 and Te29) likely represent the same individual, despite being sampled from different original material. For downstream analyses, we kept only the sample with higher sequencing depth from each pair (Ob2 and Te29).

We identified most historic Alpine ibex samples as males (12 of 14, 86%, p<0.007, Supplementary Figure S4), reflecting collectors preferences for large horns. We also identified the majority of the ancient cave samples as males (27 of 42, 64%, p<0.045, Supplementary Figure S4). While this may reflect male-biased dispersal and territory exploration in that species, it is likely also driven by collector bias since the excess males all derive from sampled full skulls, while the samples obtained from the fragmented assemblage at cave Tempiette (Ch) (Griggo et al., 2019a) showed a balanced sex ratio (10 of 24, 42%, p>0.270).

We complemented our data with published, present-day whole genomes of 29 Alpine ibex, four Iberian ibex, two Siberian ibex, two Nubian ibex (Grossen et al., 2020), Supplementary Table S1). To ensure comparability, we analyzed all available data using the ATLAS pipeline (Link et al., 2017; Marchi et al., 2022) with identical settings and relying exclusively on uncertainty-aware downstream analyses based on recalibrated genotype likelihoods. To avoid issues arising from potentially mismapped sequencing reads, particularly of the ancient samples with short reads, we i) removed reads with excessive soft-clipping or mismatches to the reference and ii) masked sites for which the cumulative depth across all ancient Alpine ibex samples was exceptionally high or low compared to the genome-wide distribution (see Methods). This resulted in 1,999 Mb on the autosomes, at which the average depth of the ancient and historic samples ranged from 0.23 X to 42.51 X (Supplementary Table S1).

Our bioinformatic treatment resulted in reliable genotype likelihoods for all present-day and ancient samples with a depth > 0.5 X, as indicated by both downsampling experiments of samples with higher depths (Supplementary Figure S5) and the fact that inferred X to autosome diversity ratios were markedly different between male and female samples (Supplementary Figure S6). In a multi-dimensional scaling (MDS) of the genetic distances between all pairs of samples, including all ibex species, all ancient samples cluster with present-day Alpine ibex samples, confirming their species designation (Supplementary Figure S7). In contrast, the data of historic samples appears to be less reliable: downsampling experiments suggested recalibration was unsuccessful for 12 out of 14 historic samples, the X to autosome diversity ratio of many historic male samples exceeded those of female samples (Supplementary Figure S6), and several historic samples fell outside the cluster of present-day Alpine ibex samples in the MDS. We suspect that these results are indicative of the harsh chemical treatment applied to these historic hunting trophies, from which DNA was extracted primarily from horns. Interestingly, two samples (SLn1 and SLn2) clearly cluster with Siberian ibex and one sample (SKm) with present-day Nubian ibex (Supplementary Figure S7A), suggesting somewhat unreliable original labelling of historic material. As a consequence, we excluded historic samples and low quality samples, and focused on the 40 ancient Alpine ibex samples with depth >0.5 X and all present-day samples.

### Stable population structure across millenia

To assess the genetic relationship among samples and investigate potential population structure, we performed a MDS on Alpine ibex samples (Figure 2A). This revealed substantial population structure among ancient Alpine ibex, a pattern confirmed by unsupervised clustering with ngsAdmix (Figure 2B, K=5), which clusters individuals by geographic region (CS, Vs, Ch). The only exceptions are the two Austrian (At) samples (SHg and THg), which fall surprisingly close to those from central Switzerland (CS). Genetic differentiation among the regional clusters was high, with F_st_ values ranging from 0.056 (between CS and At) to 0.178 (between Ch and CS) in pairwise comparisons (Figure 2C). Notably, there was also detectable geographic substructure within the CS region, for which samples clustered by cave, despite their close proximity (<95 km) and spanning ten millenia. F_st_ between the two caves with multiple samples (CSLu and CSSz) was 0.038.

**Figure 2:**
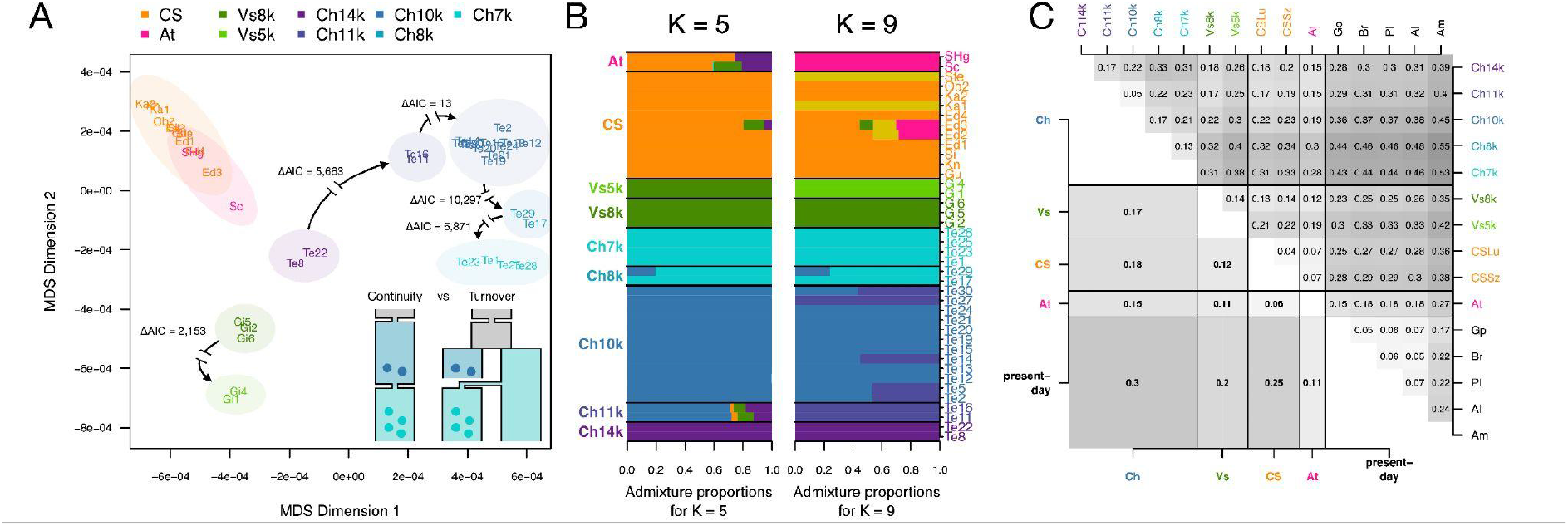
Population structure among ancient Alpine ibex populations. A) A two dimensional MDS on pairwise genetic distances. Colored ellipses indicate identified clusters. For temporally sequential clusters of the same region, continuity was assessed by contrasting the two explicit demographic models shown in the lower right: a continuity model with a potential bottleneck between the sampling times, and a turnover model with a population split that predates both clusters. Black arrows with gaps indicate cases with a significant preference for the turnover model, along with ΔAIC values for the comparison. B) Unsupervised genetic clustering with ngsAdmix for K=5 and K=9 clusters. Colors were chosen to match those used in A. C) Pairwise F_st_ matrix for all clusters (upper triangle) and regions (lower triangle). The present-day populations are ordered by their long-term effective population size as inferred by Grossen et al. (2020): Gran Paradiso (Gp), Brienzer Rothorn (Br), Pleureur (Pl), Albris (Al) and Alpi Maritime (Am).

### Ancient metapopulation dynamics

While regional population structure apparently remained stable over thousands of years, there is strong evidence for population turnover at individual locations. This is most apparent for the Ch region, within which the samples form five temporal subclusters in the MDS (Figure 2A): i) The two samples Te22 and Te8 dating to 13.2k BP and 14.2k BP, respectively, fall closest to Vs samples (Ch14k), ii) the two 11k BP samples Te11 and Te16 (Ch11k), iii) the 12 samples between 9.5-10.6k BP that fall highest on MDS2 (Ch10k), iv) the two samples Te17 and Te29 dating to 8k BP and 8.5k BP, respectively (Ch8k), and v) the four samples Te25, Te1, Te28 and Te23 < 7.5k BP (Ch7k). Two temporal subclusters are also evident for the Vs region: i) The three samples Gi2, Gi5 and Gi6 between 7.1-8.4k BP (Vs8k) fall lower on MDS2 than the two samples Gi1 and Gi4 between 4.8-5.3k BP (Vs5k). With the exception of Ch7k and Ch8k, these temporal subclusters were also identified by ngsAdmix (Supplementary Figure S8, K=5 and K=9) and most were highly differentiated (Figure 2C), including the temporally sequential clusters Ch14k and Ch11k (F_st_ = 0.17), Ch10k and Ch8k (F_st_ = 0.17) and Vs8k and Vs5k (F_st_ of 0.14). While we do not find a major effect of sampling time in the CS region, the oldest sample from Edisloch cave (Ed3, 9.5k BP) falls notably apart from the younger samples of that cave (all <8k BP) on the MDS (Figure 2A).

To test whether the observed genetic clustering by time are the result of genetic drift between discontinuous sampling or population turnovers (i.e. extinction and recolonization), we performed continuity tests by contrasting two explicit demographic models for each pair of clusters: a continuity model with a potential bottleneck between the two clusters, and a turnover in which both clusters were sampled after their populations split (Figure 1B, 2A). The turnover model was favoured in all sequential comparisons, with ΔAIC = 13 for the comparison between the Ch11k and Ch10k clusters and ΔAIC > 2152 for all others (Figure 2A, Supplementary Table S2). In contrast, for any artificial temporal split of samples of the same identified cluster, the continuity model was favoured with ΔAIC < -56 (Supplementary Figure S9), confirming the robustness of the analysis.

The combination of strong geographic population structure and regular temporal turnover at individual locations suggests that ancient Alpine ibex experienced profound metapopulation dynamics, which is well in line with their ecological niche. The species lives in alpine meadows and rock faces, today mostly at altitudes between 1500 and 3000 and avoids dense forests (Brambilla et al., 2020; Grignolio et al., 2003, 2007). This preference leads to an island-like distribution of favourable Alpine ibex habitat, at which small local populations persisted with only little gene-flow across forested valley floors.

### Near-extinction depleted species-wide but not individual diversity

To assess the impact of the near-extinction of Alpine ibex in the 19th century, we compared our ancient samples against those of available present-day samples. We included present-day samples from both the only surviving population in the Gran Paradiso area (Gp), as well as from four reintroduced populations, for which the reintroduction history and its impact on genetic diversity is well documented (Biebach & Keller, 2009; Grossen et al., 2018, 2020): Brienzer Rothorn (Br), Pleureur (Pl), and Albris (Al) from Switzerland and Alpi Marittime (Am) from Italy. Our comparison makes clear that most of the past species-wide genomic diversity of Alpine ibex was lost during the recent anthropogenic near extinction, in line with what was previously reported for mtDNA (Robin et al., 2022). This is well illustrated by an MDS of ancient and present-day samples (Figure 3A), on which present-day samples cluster tightly and outside the ancient range, a pattern confirmed by unsupervised genetic clustering with K=2 that mostly distinguishes ancient from present-day samples (Supplementary Figure S10). While we likely lack ancient samples closely representing the ancestral population of the Aosta Valley, Italy, from where all present-day samples derive (Brambilla et al., 2020), the distant placement of present-day from ancient samples is likely also driven by the extreme genetic drift the anthropogenic bottleneck imposed (Biebach & Keller, 2009; Grossen et al., 2018, 2020).

**Figure 3.**
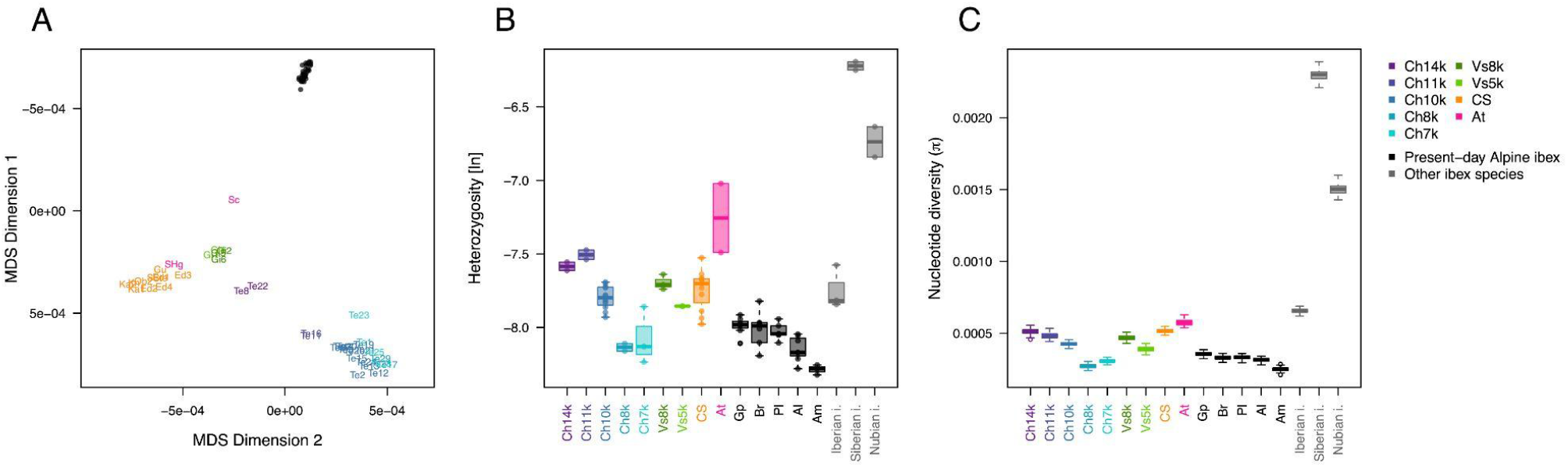
Loss of genetic diversity after near-extinction. A) A two dimensional MDS on pairwise genetic distances between all ancient and present-day Alpine ibex individuals. The Y-axis is inverted to better correspond to the MDS shown in Figure 2A. B) Individual heterozygosity estimates (dots) at autosomal putatively neutral sites, along with boxplots indicating the quantiles for each cluster. The ancient Alpine ibex populations match Figure 2. Other species for comparison include the Iberian Ibex, the Siberian Ibex and the Nubian Ibex. C) Nucleotide diversity (π) per cluster at neutral sites, along with boxplots showing the distribution across 100 bootstrap replicates. Labels as in B.

While the near extinction resulted in the loss of the majority of the species-wide genetic diversity, the diversity at the level of individuals (Figure 3B) and populations (Figure 3C) appears to have been much less severely affected than comparison to related ibex species would suggest (Grossen et al., 2020). Instead, genetic diversity was already low among ancient populations, with some populations showing lower diversity than present-day populations, particularly those affected by population turnovers (e.g. Ch8k and Vs5k, Figures 2A, 3). Median individual heterozygosity, for instance, was estimated at 4.17×10^-4^ across all ancient samples, with notable variation among both individuals and populations (Figure 3B). In contrast, median individual heterozygosity of present-day samples was estimated at 3.12×10^-4^, suggesting a loss of about 25%, which was least pronounced for the Gran Paradiso (Gp) population (median individual heterozygosity at 3.41×10^-4^, 18.2% loss) and more pronounced for reintroduced populations, particularly Albris (Al, 2.94×10^-4^, 29.4%) and Alpi Maritime (Am, 2.54×10^-4^, 39.0%), of which the latter is known to have experienced a particularly strong reintroduction bottleneck (Terrier & Rossi, 1994). Population-wide nucleotide diversity (π) similarly declined by about 40% from about 5×10^-4^ to about 3×10^-4^ (Figure 3C), but we note that estimates of ancient populations may be somewhat inflated by pooling samples spanning several thousand years. Notably, we observed similar losses of genetic diversity also after ancient population turnovers, with the turnover between the Ch10k and Ch8k, for instance, resulting in the loss of 28.9% of the median individual heterozygosity and 36.2% of the median nucleotide diversity across bootstraps. In summary, while the human-driven near extinction event of Alpine ibex led to a dramatic loss of species-wide genetic diversity, at the local scale, extinction and recolonization events resulted in some ancient populations with genetic diversity comparable to that of present-day populations.

### Accumulated mutation load in Alpine ibex

Small and isolated populations, particularly of endangered species, are known to suffer from mutation load, i.e. the accumulation of deleterious mutations, which can be seen as a proxy for genetic health (Frankham et al., 2002). For Alpine ibex, previous analyses reported increased mutation load in the reintroduced compared to the Gran Paradiso population (Grossen et al., 2020). To quantify the effect of the near-extinction event on recessive mutation load, we developed a novel, genotype-likelihood based method to estimate the fraction *f_nn_* of sites at which an individual carries two non-ancestral alleles. Applying this method to sites known to be conserved among mammals (GERP > 4) shows that present-day Alpine ibex generally carry about 3.6% more absolute recessive load than ancient individuals (median *f_nn_* of 9.67×10^-4^ vs 9.33×10^-4^), albeit with substantial individual variation and exceptionally low values for the two Austrian samples (Figure 4A). Similar to what was reported by Grossen et al. (2020), present-day populations known to have experienced particularly pronounced reintroduction bottlenecks (Al and Am) have elevated levels of recessive load (median *f_nn_* of 9.72×10^-4^ and 9.90×10^-4^, respectively). Interestingly, the ancient individuals from clusters Ch8k and Ch7k, which were likely affected by particularly strong turnover events as indicated by their low nucleotide diversity, also had elevated levels of recessive load (median *f_nn_* of 9.62×10^-4^ and 9.54×10^-4^, respectively), comparable to those of present-day populations (Figure 4A). When compared to Siberian and Nubian ibex individuals, however, all ancient individuals appear to have carried significant recessive load, as did the present-day Iberian ibex individuals (Figure 4A).

**Figure 4.**
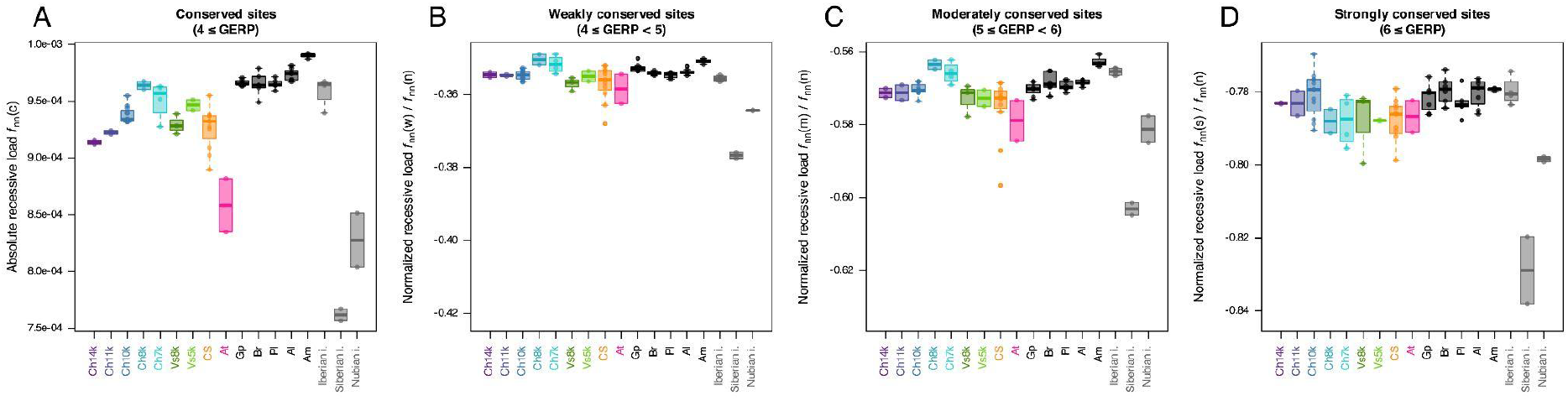
Mutation load. A) Absolute recessive mutation load quantified as the fraction f_nn_(c) of conserved sites (4 ≤ GERP) with two non-ancestral alles per individual (dot) along with boxplots indicating the quantiles per cluster or species. B) Relative mutation load quantified as f_nn_(w)/f_nn_(n), the fraction f_nn_(w) weakly conserved sites (4 ≤ GERP < 5) relative to that fraction f_nn_(n) at putatively neutral sites (GERP scores between -0.2 and 0.2). C,D) As B but for the fraction f_nn_(m) of moderately conserved and f_nn_(s) strongly conserved sites, respectively. For comparability across classes of sites, the y axes were chosen such that the lower bound is always 5/6 of the upper bound. Labels as in Figures 2 and 3.

### Evidence for purging in severely bottlenecked populations

While bottlenecks generally lead to an increased mutation load, alleles under particularly strong selection, which have been hiding in heterozygous state, may get newly exposed to selection and then removed. This process, known as purging, may result in an underrepresentation of recessive load at particularly deleterious alleles (Kirkpatrick & Jarne, 2000), an effect shown in present-day Alpine ibex for stop-gain and frameshift mutations (Grossen et al., 2020). Since the ancient data does not allow us to detect individual mutations with sufficient accuracy to be classified, we here investigated the effect of purging by contrasting *f_nn_* values obtained for sites with weak (4 ≤ GERP <5), moderate (5 ≤ GERP < 6) and strong (6 ≤ GERP) conservation scores, normalized by the *f_nn_* values obtained for neutral sites (-0.2 < GERP < 0.2).

Median normalized load across all Alpine ibex were at 0.443, 0.269 and 0.165 for weakly, moderate and strongly conserved sites, respectively, confirming that sites with high conservation scores were less variable and likely under purifying selection in Alpine ibex (Figure 4B-D). Notably, and while the normalized load of Iberian ibex was comparable, that of Siberian and Nubian ibex was considerably lower. These species do thus not only have lower levels of absolute recessive load, but also lower levels when compared to their neutral diversity. An observation, which would be in line with larger long-term population size (Robin et al., 2022) and hence overall more efficient purifying selection.

Under genetic drift alone, we expected populations that experienced stronger recent bottlenecks to show more normalized load, and that difference to be more pronounced at more conserved sites. The reason is that due to past purifying selection, the frequencies of non-ancestral alleles tend to be lower at more strongly conserved sites, allowing genetic drift to result in a stronger increase in the proportion of sites at which individuals carry two non-ancestral alleles, which we quantify using *f_nn_*. The opposite pattern of lower normalized load at more conserved sites is thus indicative of an excess removal of non-ancestral alleles in small populations, and hence purging.

In line with the drift expectation and the strong reintroduction bottlenecks, present-day Alpine ibex individuals had slightly higher normalized load compared to ancient Alpine ibex individuals across all classes of sites, with median values at 0.443, 0.270 and 0.166 for weakly, moderate and strongly conserved sites, respectively, compared to 0.442, 0.269 and 0.164 across ancient individuals. Also in line with the drift expectation, this corresponds to a relative increase of 0.1%, 0.5% and 0.9% across these classes of sites, and suggest no widespread purging in present-day Alpine ibex during the near-extinction event, at least not at the class of sites considered here.

Interestingly, however, there is some evidence for purging in populations that experienced particularly excessive drift, namely the ancient populations Ch8k and Ch7k that likely suffered from particularly strong turnover events, and the present-day Am population that suffered from the strongest reintroduction bottleneck (Terrier & Rossi, 1994). In line with the drift-expectation, the median normalized load at weakly conserved sites was 1.1% and 0.8% higher for Ch8k and Ch7k than for all other ancient individuals, and 0.7% higher for Am than for all other present-day individuals, and these values increased to 1.8%, 1.2% and 1.4% at moderately conserved sites, respectively. In contrast to the drift-expectation, however, the normalized load at strongly conserved sites was 1.0% and 0.9% lower for Ch8k and Ch7k than for all other ancient individuals, and only marginally (0.4%) higher for Am than for all other present-day individuals (Figure 4D). This excess loss of non-ancestral alleles at strongly conserved sites indicates purging: an increased efficiency of selection in these populations, despite them having experienced elevated genetic drift.

## Discussion

Since genetic diversity is the raw material for future adaptation, it is considered essential for the long-term survivability of populations and species (reviewed in Willi et al., 2022) and its protection is one of the four main goals of the CBD’s Kunming-Montreal Global Biodiversity Framework (2022). It is increasingly recognized, however, that the absolute levels of neutral genetic diversity alone is an insufficient predictor of survivability as it is strongly shaped by past demographic events and merely a proxy for the functional diversity that underlies adaptation (Abson et al., 2026; Díez-del-Molino et al., 2018; Teixeira & Huber, 2021). Instead, the impact of past population crashes should best be quantified in terms of the loss in both neutral and functional genetic diversity, and the accumulation of deleterious mutation load, ideally through the direct comparison of populations through time (Díez-del-Molino et al., 2018; Willi et al., 2022). While the usefulness of time-series data is well recognized in conservation genetics (Epstein et al., 2016; Thalmann et al., 2011; Węcek et al., 2016), systematic comparison of genome-wide diversity of endangered species remains limited to only shallow evolutionary times (Dussex et al., 2021; Lehnert et al., 2019) or few samples (Lucena-Perez et al., 2024).

Here we present ancient whole genomes for 40 Alpine ibex from across the entire former range and spanning the last 14 ky. They make clear that Alpine ibex existed for thousands of years as a highly dynamic metapopulation that consisted of small populations, connected by limited gene flow only, which regularly experienced local extinction and recolonization. This is consistent with their narrow ecological niche of alpine meadows and rocky cliffs, and their avoidance of forests (Grignolio et al., 2003, 2007). With the retreat of the glaciers at the end of the last glacial cycle, and the change of the dominant lowland vegetation type to forests (Tinner & Kaltenrieder, 2005), Alpine ibex have been restricted to the narrow and dynamic band between the treeline at the lower end and mountain tops or retreating glaciers at the upper end, with valleys functioning as effective barriers. This habitat remained dynamic and affected by small climatic fluctuations with higher areas becoming arid or glaciated during cold phases and lower mountain tops becoming forested during warm phases. Such fluctuations, along with chance events, frequently resulted in extinction of the local population, followed by recolonization when the habitat improved again, as is well documented in our data at both the Tempiette cave and the Giétroz cave. Notably, most of the detected turnover events also resulted in a significant loss of genetic diversity, suggesting a depletion of the local gene pool and the eventual regional extinction, as evidenced by the absence of Alpine ibex fossils after 7100 BP at the Tempiette cave (Griggo et al., 2019a and this study) and between 4800 BP and 1800 BP at the Giétroz cave (Reynaud Savioz & Blant, 2021), despite the presence of fossils of other species.

In line with their narrow niche and island-like geographic distribution, as well as indications from ancient mtDNA (Robin et al., 2022), we here report low genetic diversity of past Alpine ibex populations long before anthropogenic influences became dominant in the Alps (Boxleitner et al., 2017). The neutral diversity of ancient Alpine ibex populations was significantly lower than that of present-day Nubian and Siberian ibex, and comparable to that of the formerly bottlenecked Iberian ibex. Similarly, ancient Alpine ibex populations carried substantial recessive mutation load, similar to that of the Iberian ibex, but much higher than those of the Nubian and Siberian ibex, reflecting their much larger population sizes. Nonetheless, the anthropogenic near-extinction event is detectable with present-day Alpine ibex populations having about 20% less neutral genetic diversity at the individual and about 40% less at the population level than ancient populations, and a slight (3%) increase in recessive mutation load. As a result, the diversity of present-day populations is on par with that of the last populations at the Tempiette and Giétroz caves.

The observed small difference between ancient and present-day, reintroduced populations regarding their genetic makeup may have contributed to the rapid recovery of Alpine ibex. Selection is generally considered less efficient in small populations, allowing highly deleterious recessive alleles to segregate at low frequency. After an episode of rapid drift, such as after a severe near-extinction event, many of those variants may have been lost. But variants that survived likely segregated at frequencies high enough to regularly be homozygous and thereby considerably reducing their carriers fitness, a phenomenon known as inbreeding depression. In line with the weak selection expectations, we found higher mutation load in Alpin ibex compared to other species. We propose, however, that the regular episodes of local bottlenecks over thousands of years likely led to an efficient purging of the most deleterious alleles in Alpine ibex, keeping the inbreeding depression in reintroduced populations at detectable (Bozzuto et al., 2019; Brambilla et al., 2015, 2018) but largely mild levels.

To quantify mutation load, we relied on GERP scores that reflect the conservation of a site across mammals. These scores appear to generally be good proxies for the deleteriousness of derived alleles in Alpine ibex, evidenced by the much lower load at highly compared to weakly conserved sites. There is nonetheless likely considerable variation in the deleteriousness of derived alleles even among sites with similar conservation scores (Huber et al., 2020), and the excess load of Alpine ibex may be mostly composed of sites highly conserved across mammals in general, but with milder effect in Alpine ibex. In line with this interpretation, we do not find evidence for widespread additional purging during the near-extinction event, except for the population that experienced the most severe reintroduction bottleneck, for which the additional purging appears on par with what we found in the ancient populations that experienced the most severe recolonization bottlenecks.

While the genetic diversity of present-day populations may not differ much from that of ancient populations, the loss of genetic diversity was dramatic at the species level. Due to their strong geographic structure, most of the genetic diversity of ancient Alpine ibex existed not within but between populations, as illustrated by high F_st_ values even between populations in close proximity, and remained geographically restricted for thousands of years. During the anthropogenic near-extinction event, all of this diversity was lost, with the exception of the small fraction that segregated within the only surviving population in the Gran Paradiso area. While species-level diversity may be much less relevant than mutation load for the short-term recovery, it may be essential for long term adaptation. Since all extant populations share a very similar genetic makeup, future environmental changes, including new diseases, may prove fatal for current Alpine ibex populations as no reservoir of novel genetic diversity exists to replenish areas affected by local extinctions. While future scenarios remain challenging to predict, the time-series reported here, particularly from the Tempiette and Giétroz caves, serve as reminders of both the relevance of a healthy genetic pool for successful recolonization, as well as the very real possibility of at least regional extinctions.

## Methods

### Sampling and laboratory protocols

#### Sampling and DNA extraction

We screened a total of five Iberian ibex and 88 putative Alpine ibex samples, including ancient samples from caves and glaciers, as well as historic samples, mostly hunting trophies (Supplementary Table S3). For the ancient samples originating from caves, we extracted aDNA mainly from petrous bones and teeth. For the historic samples, mostly horns were used. Before sample extraction, samples were irradiated with UV for 30 min and surfaces were cleaned with 7% Sodium hypochlorite/NaClO. The outermost part of the bone or horn was removed with a dental drill and tungsten steel drill bits (Alpine Orthodontics, H1-014-HP) before extracting the bone powder under very low rotation rate to avoid further DNA denaturation due to heat. For the historic hunting trophies we were not able to perform the sampling in the laboratory and worked with a portable precision drill (Dremel 9100-21 Fortiflex) and tungsten steel drill bits (Alpine Orthodontics, H1-014-HP) as described in (Sirak et al., 2017). Here, we changed the drill bit after removing the most outer part of the horn surface and extracted approximately 100 - 200 mg of bone powder into an Eppendorf tube. This allowed a minimally invasive sampling of the valuable historic specimens on spot.

DNA extraction was performed as described in Robin et al., 2022. In short, The QIAquick® PCR Purification Kit was used to extract DNA from the bone powder implementing the following modifications of the standard manufacturers protocol: We used 100 mg bone powder and added 50 µl Proteinase K (800 units/ml, New England Biolabs Inc.) and 950 µl EDTA. The lysis was conducted at 37 °C overnight and the overnight digestion was centrifuged at 14000 rpm for 5 minutes in a tabletop centrifuge, mixed with 10 ml binding buffer (without tween) and 400ul 3M sodium acetate. We centrifuged the resulting suspension at 1500 rpm for 15 minutes. We stored the eluted samples at 4 °C.

#### Library preparation and sequencing

We built double-stranded DNA libraries for all samples as described in Robin et al., 2022 and took the following strict precautions to avoid contamination. During library construction, all pre-amplification steps were performed in a specialised aDNA clean laboratory at the Institute for Evolutionary Medicine, University of Zürich. Non-template controls were included for all runs and library construction followed the protocol described in Kircher et al., 2012 and Meyer & Kircher, 2010. To produce high coverage we first equilibrated all libraries to 10 nM using a TapeStation 2200 with D1000 ScreenTape® (5067-5582) with D1000 Reagents (5067-5583).

We performed an initial screening run on an Illumina NextSeq-500 in mid-output mode with 150 cycles (2*75+8+8) at the Functional Genomic Centre Zürich (FGCZ) to assess the state of DNA preservation and endogenous DNA content. The most promising samples, generally those with >30% high endogenous DNA content after removing duplicate reads (Supplementary Table S3), were repooled based on per sample depth achieved and subsequently sequenced on an Illumina NovaSeq 6000 using the NovaSeq Xp workflow and S4 v1.5 flow cells (2 x 100 bp) at the facility of the National Genomics Infrastructure (NGI), Sweden. Each sample was sequenced on two to four independent flow cells to reduce the influence of cluster specific sequencing errors and hence, batch effects.

#### Previously published samples

We included published genomes of present-day Bezoar (N=3), domestic sheep (N=2) and domestic goat (N=6) from (Alberto et al., 2018) and of Iberian ibex (N=4), Siberian ibex (N=2), Nubian ibex (N=2), Makhor (N=1) Alpine ibex (N=29) from (Grossen et al. 2020). We grouped Alpine ibex individuals into five populations: the surviving population in the Gran Paradiso area (Gp), the reintroduced population Alpi Marittime (Am), and the initial three reintroduced populations, including sequenced individuals from populations that derived from these: we treat the three individuals from each of Brienzer Rothorn and Bire Öschinen collectively as “Brienzer Rothorn (Br)”, the three individuals from each of Pleureur and Weisshorn collectively as “Pleurer (Pl)” and the three individuals from Albris, the three individuals from Oberbauenstock and the single individual from Rheinwald collectively as “Albris (Al)”. We used published genomes of present-day Bezoar (N=3), domestic sheep (N=2) and domestic goat (N=6) (Alberto et al., 2018) downloaded from ENA (project PRJEB7436) as outgroups.

#### Dating

We subjected all ancient samples used for whole genome sequencing to AMS-14C dating at the Laboratory of Ion Beam Physics, ETH Zurich in Switzerland, unless they were previously dated (Supplementary Table S1) (Griggo et al., 2019a; Reynaud Savioz et al., 2018; Reynaud Savioz & Blant, 2021). Raw radiocarbon dates were recalibrated using the R package rcarbon (Crema & Bevan, 2021) using the intcal20 recalibration curve (Reimer et al., 2020). The age and sampling location of historic samples was determined based on museum entries, oral statements or from historical records and can therefore be imprecise. Hereafter, we follow the suggestion of Raxworthy & Smith, 2021 and refer to samples older than 300 years as ancient, to those younger than 300 years as historic, unless they lived in the last few years, in which case we refer to them as present-day.

### Bioinformatic processing

#### Raw data analysis

The raw data was processed with ATLAS (commit 7b82b51, obtained from https://bitbucket.org/wegmannlab/atlas) (Link et al., 2017), using the ATLAS-pipeline (commit dc411a5, obtained from https://bitbucket.org/wegmannlab/atlas-pipeline) as described in (Marchi et al., 2022), with minor modifications. In short, we used the Gaia module of the ATLAS-Pipeline to 1) check sequencing quality with fastqc 0.11.8 (www.bioinformatics.babraham.ac.uk/projects/fastqc/), 2) trim raw reads with Trim Galore 0.6.7 (https://github.com/FelixKrueger/TrimGalore) with no quality filter, a length filter of 30 bp and option --retain unpaired, 3) verify trimming and the quality of the remaining sequences again with fastqc, 4) align reads to the Alpine ibex genome CapIbe1.0 (F. G. Çilingir et al., 2025) using bwa 0.7.17 -mem (Li, 2013) with options -t 10 -M, 5) filter out reads with mapping quality <30 using SAMtools 1.9 (Danecek et al., 2021) with options -q30, 6) add informative read groups with picard-tools 2.26.11 (http://broadinstitute.github.io/picard/) AddOrReplaceReadGroups, 7) sort and index BAM files with SAMtools index and sort, and 8) obtain statistics such as read counts, read lengths, sequencing depth and transition and transversion rates with ATLAS using the tasks BAMDiagnostics and refDiffTable.

The second ATLAS-pipeline module Rhea was then used to perform local realignment around insertion and deletion (In-Del) positions with GATK version 3.8 (DePristo et al., 2011). For a computationally efficient and parallelizable approach, we followed a two-step procedure similar to that initially proposed by Marchi et al., 2022., in which local realignment was performed on each sample individually, but informed by a subset of samples representing the full species diversity (the guidance set). Here, we obtained the guidance set from 34 samples (Supplementary Table S1) by first identifying potential target intervals using GATK RealignerTargetCreator and then downsampling their data to 5 X at identified target sites and to 0.1 X throughout the rest of the genome using the ATLAS task downsample with option --downsampleTarget. We then performed local realignment with GATK IndelRealigner on each sample individually on the target intervals of the guidance set, while also providing the downsampled reads of the guidance set.

#### Sex identification

For all ancient and historic Alpine ibex samples, we used the task BAMDiagnostics in ATLAS with option --perChromosome to count the number of reads that aligned to each scaffold, ignoring reads with a mapping quality below 30 (--filterMQ [30,]) and fragments with a length smaller than 40 (--filterFragmentLength [40,]). We then used BeXY (Caduff et al., 2024) to infer sex karyotypes and sex-linked scaffolds using the task infer and setting a minimal scaffold length filter of 10^5^.

#### Filtering

To mitigate potential mapping issues, we identified and masked positions that had either extraordinarily high or low sequencing depth across all ancient samples, or positions on the X chromosome for which the depth ratio between male and female samples differed from the chromosome-wide expectation.

Using the ATLAS-pipeline module DepthFilter, which first ran the ATLAS task pileup to obtain the histogram of the per-site cumulative depth across all samples, and then the ATLAS task pileupToBed with option --quantile 0.99 to fit a negative binomial distribution to this histogram, identify the 0.5% and 99.5% quantiles of that distribution, and write a bed file containing all positions for which the cumulative depth was inside that range for downstream use. These steps were repeated for males and females separately on the X-chromosome (scaffold_2).

To identify positions on the X-chromosome at which the proportion of cumulative depth of male samples diverge from chromosome-wide expectations, we implemented a novel, iterative approach as a new option to the task pileupToBed in ATLAS. In each iteration, it first determines the fraction *m* of the male cumulative depth out of the combined depth across all sites considered. It then performs a two-sided binomial exact test with fraction *m* on the cumulative depth at each site and identifies outlier sites as those with p-value < 0.001, considering all remaining sites in the next iteration. It then repeats these steps until the fraction *m* no longer changes, and writes a bed file containing the considered sites for downstream use.

For the X chromosome, we only considered sites that passed all of the three filters by intersecting the corresponding bed files with ATLAS task bed. The final bed files were then filtered to only contain genomic regions of at least 100 consecutive loci, which together covered 82.79% of the reference.

#### Genotype likelihoods

The BAM files obtained with the Rhea module were processed with the ATLAS-pipeline modules Perses and Pallas to 1) merge paired-end reads with task mergeOverlappingReads and options --filterSoftClips 0.0,0.1 --filterFragmentMismatches 0.0,0.05, 2) to estimate the likelihood of sequencing errors and PMD with task estimateErrors using options --ploidy 1 on 10Mb ultraconserved Genomic Evolutionary Rate Profiling (GERP; Cooper et al., 2005) scored sites obtained from (G. Çilingir et al., 2025) 3) to compute genotype likelihoods with task GLF while providing updated error rates with option --RGInfo and the depth filter with --regions.

### Population genomic analyses

#### Relatedness

We assessed relatedness between pairs of samples using NgsRelate v2.3.0 (Korneliussen & Moltke, 2015). As an input we provided the VCF file as described above. Per default NgsRelate will estimate the allele frequencies using the individuals in the provided VCF. We instead estimated allele frequencies using the ATLAS task alleleFreq and annotated the VCF using BCFtools version 1.12 (Narasimhan et al., 2016) to add the alternate allele frequencies to the INFO field.

#### Ancestral states

We obtained ancestral state information on ibex in three steps: First, we obtained bi-allelic genotype likelihoods with ATLAS task majorMinor from the GLF files of seven representatives of the Markhor, Bezoar, domestic goat and domestic sheep, which we considered as outgroup species in our analysis (Supplementary Table S1), using the options --method Skotte --minMAF 0 and --minSamplesWithData 7. Second, we estimated allele counts for each site with ATLAS task alleleCounts using the option --outFormat withAlleles. Third, we identified the ancestral state with ATLAS task ancestralAlleles using options --minorCountMaximum 0 and --totalCountMinimum 14 to limit to sites at which all samples had data (i.e. all 14 alleles were observed) and all samples shared the same allele.

#### Subset of neutral and conserved sites

We conducted population genomic analyses on two subsets of sites based on their mammalian conservation scores (GERP) obtained from G. Çilingir et al., 2025 and restricted to sites for which ancestral states could be inferred: a “neutral” set consisting of sites with -0.2 <= GERP < -0.2 (208,433,357 loci) and a “conserved” set consisting of all sites with GERP >= 4 (20,585,823 loci). The allele identified as ancestral was added to every locus using bedtools version 2.3 (Quinlan & Hall, 2010) getfasta with option –bedOut. To increase computation speed, we recreated individual genotype likelihood files using the ATLAS task GLF for both subsets using option --regions. For some analyses, we further subdivided the conserved set into “weakly” (4 ≤ GERP <5, 11,766,766 loci), “moderately” (5 ≤ GERP < 6, 6,641,598 loci) and “strongly” (6 ≤ GERP, 2,177,459 loci) conserved subsets.

#### Population structure

We estimated genetic distances at all neutral loci between all pairs of ibex samples using the ATLAS task geneticDist. These distances were visualized using multidimensional scaling performed in R version 4.5.3 (R Core Team, 2025).

We further performed admixture analysis with ngsAdmix (Skotte et al., 2013) on all 40 ancient Alpine ibex samples, as well as on all 79 present-day and ancient Alpine ibex samples. To do so, we identified the major and minor alleles at all neutral loci and encoded bi-alleleic genotype likelihoods in a VCF file with ATLAS task majorMinor and options --method Skotte, --minSamplesWithData N/2 and --minMAF 0.05, where N is the number of samples in the data set. VCF files were converted into beagle format using ATLAS convertVCF. We ran ngsAdmix for K values from 2 to 12, with 20 repetitions for each K-value and option --maxIter 4000.

#### Genetic diversity

We inferred individual heterozygosity using the ATLAS task summaryStats and option –estimators HKY85. To infer population-specific nucleotide diversity (π) (Tajima, 1989), we first created population-specific saf files at neutral sites using the ATLAS task saf and the option --fasta to provide the ancestral states, and then used winSFS 0.7.0 (Rasmussen et al., 2022) with default options to infer unfolded one-dimensional SFS. We calculated the nucleotide diversity for each population per autosome with the tools realSFS saf2theta followed by thetaStat do_stat, which are both supplied in ANGSD version 0.941-22-gc877e7f (Korneliussen et al., 2013, 2014). We then calculated π from the resulting .pestPG files as the sum of pairwise theta (nucleotide diversity) divided by the number of sites. The analysis was bootstrapped with 100 replicates using 10 Mb neutral loci each.

#### Mutation load

We estimated recessive mutation load as the fraction *f_nn_(x)* among a subset x of genomic positions at which an individual carries two non-ancestral alleles. We estimated this fraction directly from the full set of genotype likelihoods encoded in GLF files, for which we implemented the new task mutationLoad in ATLAS (see supplementary information for details on the method). We ran this task for the conserved subsets of sites defined above and provided ancestral allele with option --alleles. We further calculated the normalized recessive load as their fraction *f_nn_(x)* divided by the fraction *f_nn_(n)* obtained for putatively neutral sites.

#### Demographic inference

We created population-specific saf files at neutral sites using the ATLAS task saf, then used winSFS 0.7.0 (Rasmussen et al., 2022) with default options to infer folded two-dimensional SFS. We used fastsimcoal2 2.7093 (Excoffier et al., 2021) to infer demographic parameters under a bottleneck model and a population turnover model. We modeled each sample according to its maximum a posteriori calibrated date while assuming a generation time of 8 years and used poolSFS to simulate their collective SFS (Marchi et al., 2024). For each model, we inferred all parameters jointly (population sizes and split times, as well as bottleneck times and sizes, if applicable) by maximizing the model likelihood in 1000 independent runs with 50 expectation conditional maximisation (ECM) cycles per run and 1 x 10^6^ coalescent simulations to estimate the expected SFS. The command line used for the estimation was:

fsc -t model.tpl -n1000000 -d -e model.est -M -L50 --multiSFS

Because the polymorphic sites in our data are not strictly independent, the computed likelihoods are composite likelihoods. The associated ΔAIC (ΔAIC = (2*5 - 2*LL_bottleneck_) - (2*6 - 2*LL_turnover_)) might therefore be overestimated (Marchi et al., 2024).

## Data availability

Raw sequencing data (FASTQ-files) of ancient whole genome data produced for this study have been deposited to the European Nucleotide Archive (ENA: PRJEB84031) and are publicly available as of the date of publication. The newly developed ATLAS features are available through the ATLAS git repository at https://bitbucket.org/wegmannlab/atlas.

## Acknowledgements

We would like to acknowledge the different institutions for granting us access to ancient and historical Alpine ibex samples. The following institutions supplied samples for the aDNA analysis: Archäologischer Dienst Graubünden, Naturhistorisches Museum Bern, Nationalpark Hohe Tauern, Naturhistorisches Museum Winterthur, Natur-Museum Luzern, Stiftung Naturerbe Karst und Höhlen Obwalden, Naturhistorisches Museum Basel, Naturhistorisches Museum Freiburg, Naturmuseum Olten, Zoologisches Museum Zürich, Institut Suisse de Spéléologie et de Karstologie ISSKA, Université Grenoble Alpes, Université Toulouse, Université Neuchâtel, Stift Kremsmünster, Dommuseum Salzburg, Verein Haus der Natur - Museum für Natur und Technik, Erzabtei St. Peter, Salzburg, Landesmuseum für Kärnten, Oberösterreichisches Landesmuseum Linz, Österreichische Paläontologische Gesellschaft, Universalmuseum Joanneum and Abteilung für Provinzialrömische Archäologie und Feldforschung, Apothekenmuseum Winkler, Innsbruck, Museum Schloss Moosham. We furthermore received samples from private owners and hence like to thank Fam. Voithofer (Gasthof Senningerbräu, Bramberg am Wildkogel), Fam. Lankmayer (Zehenthof, Göriach), Fam. Kolbitsch (Hotel Bräu, Zell am Ziller). We are grateful to Werner Müller for his contribution to the collection and curation of the ancient samples. We thank Iris Biebach, Glauco Camenisch and Lukas Keller for providing early access to some of their data for internal validation and for insightful discussions regarding the analyses. We acknowledge the Functional genomics centre Zurich for sequencing and the National Genomics Infrastructure in Stockholm funded by the Science for Life Laboratory, the Knut and Alice Wallenberg Foundation and the Swedish Research Council, and SNIC/Uppsala Multidisciplinary Centre for Advanced Computational Science for assistance with massively parallel sequencing and access to the UPPMAX computational infrastructure. This work was supported by the University of Zürich through a University Research Priority Program “Evolution in Action” pilot project grant, the Swiss National Science Foundation (31003A_182343 to C.G. and 310030_200420 to D.W.) and by FORMAS (grant numbers 2018-01640 and 2015-676 to L.D.). All computations were performed at the IBU cluster of the University of Bern.

