## Supplementary information and Figures for "Paleogenomic time series reveal past population dynamics and their relevance for the conservation of the Alpine ibex"

### Supplementary Methods for Mutation Load

Here we are concerned with quantifying the mutation load of individuals, which is usually obtained by simply counting the number of sites at which and individual is heterozygous or homozygous for a deleterious variant for sites classified as potentially deleterious *a priori*, either based on conservation scores or effect predictions. The reliance on accurately called genotypes, however, limits this analysis to high quality genomes or a small set of highly ascertained sites, which are unlikely representative of the genome-wide load and difficult to compare across populations. To extend the inference of mutation load to genome data of lower quality, as is common for ancient samples, the uncertainty of genotypes must be accounted for. Here, we develop an appproach to infer mutation load from genotype likelihoods: the probability of observing the sequencing data given the underlying true genotype and sequencing error rates.

Consider a set of sites $l = 1, \dots, L$ and let $g_l = AA, AC, AG, \dots, TT$ denote the genotype at site $l$. We assume that at each site a "preferred" allele $r_l = A, C, G, T$ is known and seek to infer the frequency of each of four genotype classes:

1. Homozygous for the preferred allele ($g_l = r_l r_l$), denoted by $rr$.
2. Heterozygous for the preferred and an alterative allele ($g_L = r_l a, a \neq r_l$), denoted by $ra$.
3. Homozygous for an alterative allele ($g_l = aa, a \neq r_l$), denoted by $aa$.
4. Heterozygous for two alternative alleles ($g_l = ab, a, b \neq r_l, a \neq b$), denoted by $ab$.

Let $\mathcal{C} = \{rr, ra, aa, ab\}$ denote the set of all possible classes and $\boldsymbol{\pi} = (\pi_{rr}, \pi_{ra}, \pi_{aa}, \pi_{ab})$ the vector of frequencies of these class with $\sum_{c\in\mathcal{C}} \pi_c = 1$. Let us further denote by $|c|$ the sizes of class $c \in \mathcal{C}$, which is the number of possible genotypes in that class with $|rr| = 1$ and $|ra| = |aa| = |ab| = 3$. Finally, let us assume that the possible genotypes $g \in c$ are equally distributed within class $c$. The likelihood of full data $\boldsymbol{d} = (d_1, \dots, d_L)$ is then given by

$$\mathcal{L}(\boldsymbol{\pi}) = \mathbb{P}(\boldsymbol{d}|\boldsymbol{\pi}, \boldsymbol{r}) \quad = \quad \prod_l \sum_g \mathbb{P}(d_l|g)\mathbb{P}(g|\boldsymbol{\pi}, r_l),$$

where $\boldsymbol{r} = (r_1, \dots, r_L)$, the $\mathbb{P}(d_l|g)$ are genotype likelihoods and for a genotype $g$ belonging to class $c$,

$$\mathbb{P}(g|\boldsymbol{\pi}, r_l) = \frac{\pi_c}{|c|} = \begin{cases} \pi_{rr} & \text{if } g = r_l r_l \\ \frac{\pi_{ra}}{3} & \text{if } g = r_l a; a \neq r_l \\ \frac{\pi_{aa}}{3} & \text{if } g = aa; a \neq r_l \\ \frac{\pi_{ab}}{3} & \text{if } g = ab; a, b \neq r_l; a \neq b \end{cases}$$

We infer the parameters $\boldsymbol{\pi}$ using an EM algorithm.

**E-step** Let $\boldsymbol{g} = (g_1, \dots, g_L)$ denote the vector of genotypes at locations $l = 1, \dots, L$. The complete data likelihood is given by

$$\mathcal{L}_c(\boldsymbol{\pi}) = \mathbb{P}(\boldsymbol{d}, \boldsymbol{g}|\boldsymbol{\pi}, \boldsymbol{r}) = \prod_l \mathbb{P}(d_l|g_l)\mathbb{P}(g_l|\boldsymbol{\pi}, r_l),$$

the complete data log-likelihood by

$$\ell_c(\boldsymbol{\pi}) = \log \mathbb{P}(\boldsymbol{d}, \boldsymbol{g}|\boldsymbol{\pi}, \boldsymbol{r}) = \sum_l \left[\log \mathbb{P}(d_l|g_l) + \log \mathbb{P}(g_l|\boldsymbol{\pi}, r_l)\right]$$

and the expected complete data log-likelihood by

$$Q(\boldsymbol{\pi}; \boldsymbol{\pi}') = \mathbb{E}[\ell_c(\boldsymbol{\pi})|\boldsymbol{d}; \boldsymbol{\pi}'] = \sum_l \sum_g \left[\log \mathbb{P}(d_l|g) + \log \mathbb{P}(g|\boldsymbol{\pi}, \boldsymbol{r})\right] \mathbb{P}(g|d_l, \boldsymbol{\pi}', \boldsymbol{r}),$$

where

$$\mathbb{P}(g|d_l, \boldsymbol{\pi}', \boldsymbol{r}) = \frac{\mathbb{P}(d_l|g)\mathbb{P}(g|\boldsymbol{\pi}', \boldsymbol{r})}{\sum_\gamma \mathbb{P}(d_l|\gamma)\mathbb{P}(\gamma|\boldsymbol{\pi}', \boldsymbol{r})}$$

Only the second part $Q_2$ of this sum depends on $\boldsymbol{\pi}$. We have

$$\begin{aligned} Q_2(\boldsymbol{\pi}; \boldsymbol{\pi}') &= \sum_l \sum_g \log \mathbb{P}(g|\boldsymbol{\pi}, \boldsymbol{r})\mathbb{P}(g|d_l, \boldsymbol{\pi}', r_l) \\ &= \sum_{c \in \mathcal{C}} \log \frac{\pi_c}{|c|} \sum_l \sum_{g \in c} \mathbb{P}(g|d_l, \boldsymbol{\pi}', r_l). \end{aligned}$$

**M-step** We have to maximize $Q_2$ subject to the constraint $\sum_{c \in \mathcal{C}} \pi_c = 1$. For this reason we form the Lagrangian

$$\mathrm{L}(\boldsymbol{\pi}, \mu) = Q_2(\boldsymbol{\pi}; \boldsymbol{\pi}') - \mu(\sum_{c \in \mathcal{C}} \pi_c),$$

where $\mu$ is the Lagrangian multiplier whose partial derivative set to 0 is

$$\frac{\partial}{\partial \mu} \mathrm{L}(\boldsymbol{\pi}, \mu) = \sum_{c \in \mathcal{C}} \pi_c - 1 = 0.$$

The partial derivatives of $\boldsymbol{\pi}$ are

$$\frac{\partial}{\partial \pi_c} \mathrm{L}(\boldsymbol{\pi}, \mu) = \frac{1}{\pi_c} \sum_l \sum_{g \in c} \mathbb{P}(g|d_l, \boldsymbol{\pi}', r_l) - \mu,$$

from which we get the estimates

$$\hat{\pi}_c = \frac{\sum_l \sum_{g \in c} \mathbb{P}(g|d_l, \boldsymbol{\pi}', r_l)}{\sum_l \sum_g \mathbb{P}(g|d_l, \boldsymbol{\pi}', r_l)}.$$

Once the estimates of $\boldsymbol{\pi}$ are obtained, they may be used to calculate mutation load:

- Additive load: the sum $\frac{1}{2}\pi_{ra} + \pi_{aa} + \pi_{ab}$ indicative of load under an additive fitness model under which a heterozygous genotype contributes have as much to the load as a homozgyous deleterious genotype.
- Dominant load: the sum $\pi_{ra} + \pi_{aa} + \pi_{ab}$ is indicative of load under a dominant fitness model where already a single deleterious allele contributes fully towards load.
- Recessive load: The sum $\pi_{aa} + \pi_{ab}$ is indicative of load under a recessive fitness model under which two non-preferred alleles are required to confer negative fitness effects.

In the main text, we assume the ancestral state is the preferred allele and refer to the sum $f_{nn} = \pi_{aa} + \pi_{ab}$ as the fraction of sites at which an individual carries two non-preferred alleles. We then compute this fraction for different classes of sites and interpret this as a measure of recessive load whenever using a set of conserved sites.

### Supplementary Figures

Figure S1: Recalibration of raw radiocarbon dates, performed with the R package rcarbon (Crema and Bevan, 2021) and using the intcal20 recalibration curve (Reimer et al., 2020). The raw radiocarbon dates are indicated on the Y-axis. The solid black line indicates the maximum a posteriori (MAP) recalibrated date, the dashed lines indicate the minimum and maximum of all given 95.4% ranges combined.

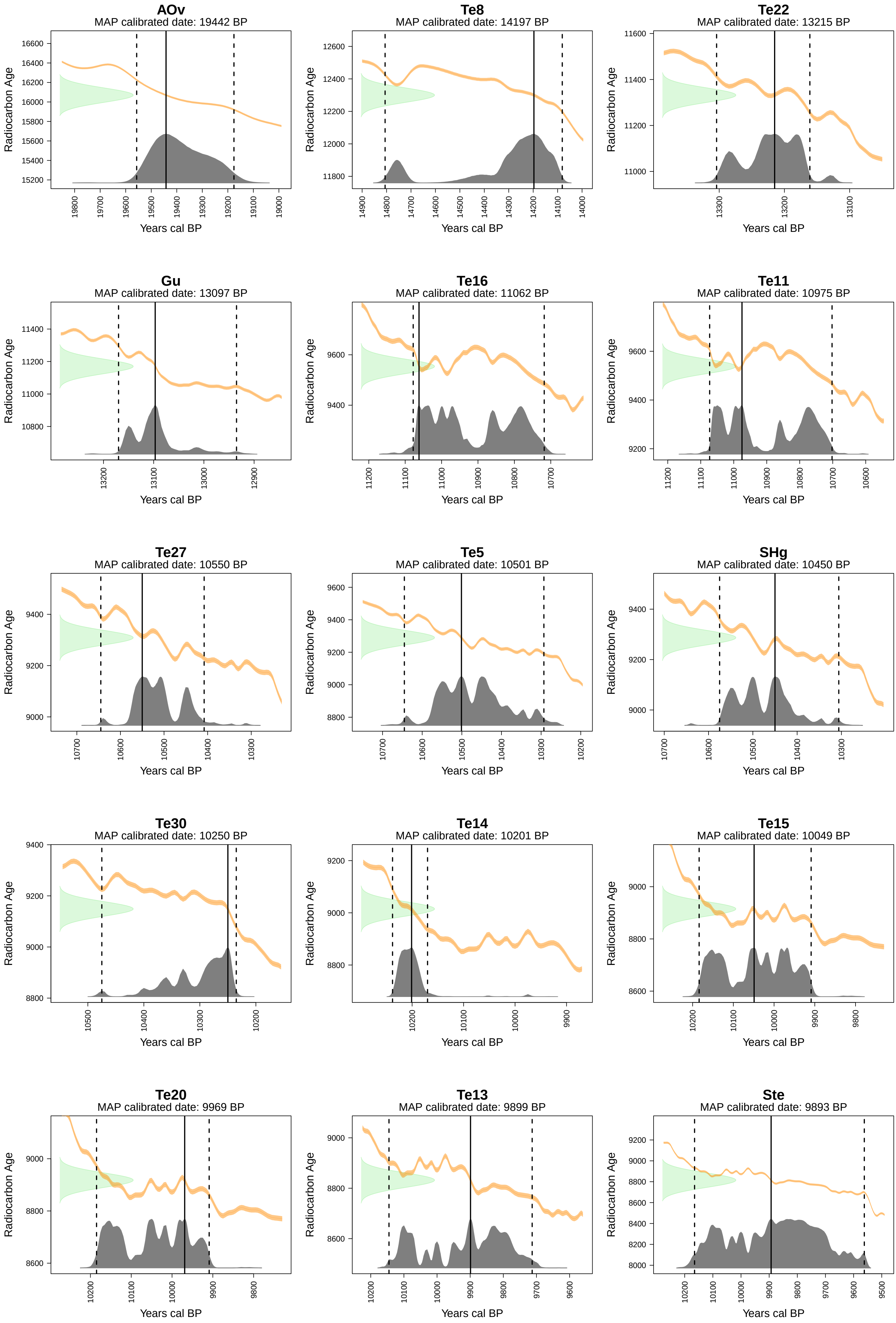
AOv
MAP calibrated date: 19442 BP
Te8
MAP calibrated date: 14197 BP
Te22
MAP calibrated date: 13215 BP
Gu
MAP calibrated date: 13097 BP
Te16
MAP calibrated date: 11062 BP
Te11
MAP calibrated date: 10975 BP
Te27
MAP calibrated date: 10550 BP
Te5
MAP calibrated date: 10501 BP
SHg
MAP calibrated date: 10450 BP
Te30
MAP calibrated date: 10250 BP
Te14
MAP calibrated date: 10201 BP
Te15
MAP calibrated date: 10049 BP
Te20
MAP calibrated date: 9969 BP
Te13
MAP calibrated date: 9899 BP
Ste
MAP calibrated date: 9893 BP
Radiocarbon Age
Years cal BP

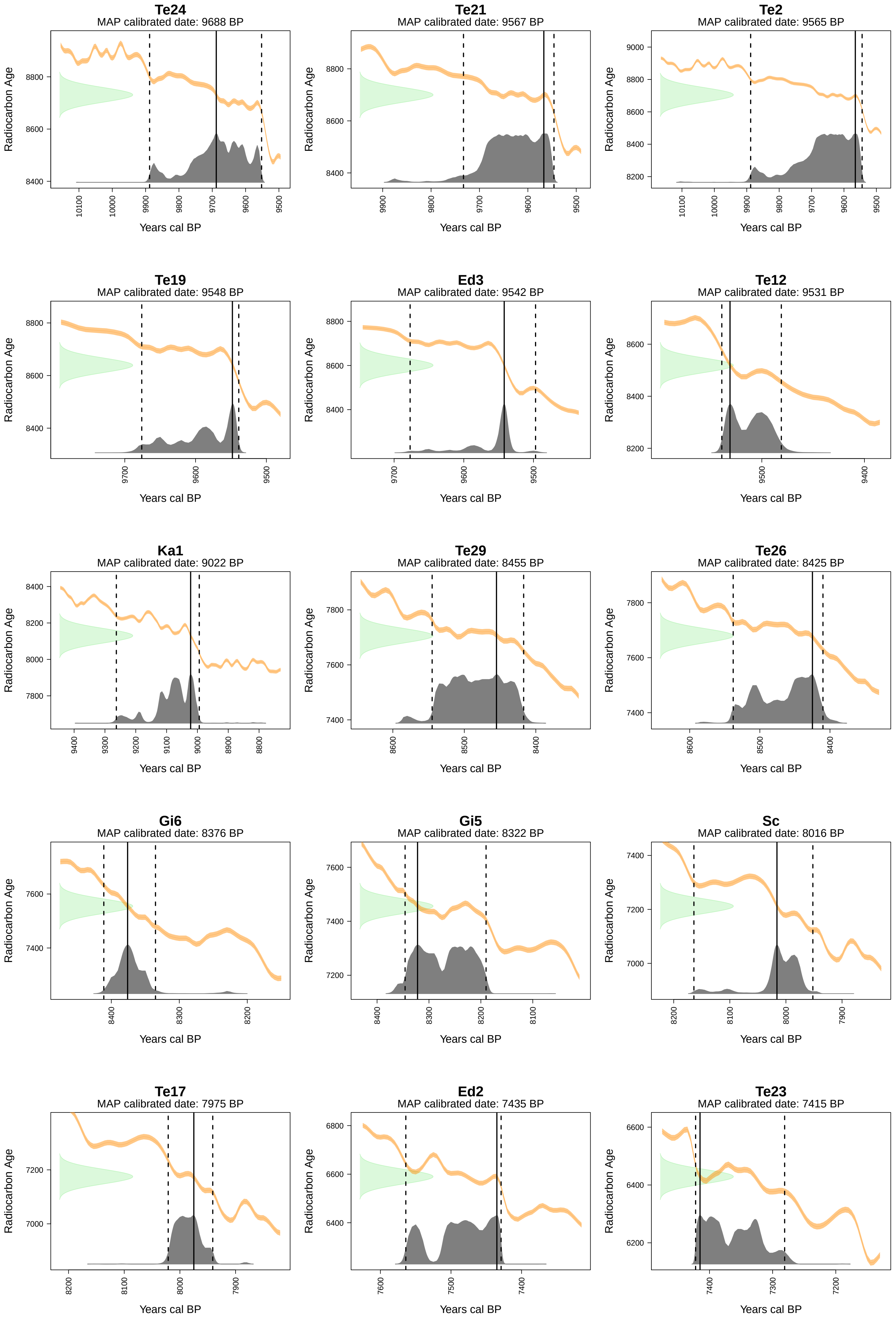
Te24
MAP calibrated date: 9688 BP
Te21
MAP calibrated date: 9567 BP
Te2
MAP calibrated date: 9565 BP
Te19
MAP calibrated date: 9548 BP
Ed3
MAP calibrated date: 9542 BP
Te12
MAP calibrated date: 9531 BP
Ka1
MAP calibrated date: 9022 BP
Te29
MAP calibrated date: 8455 BP
Te26
MAP calibrated date: 8425 BP
Gi6
MAP calibrated date: 8376 BP
Gi5
MAP calibrated date: 8322 BP
Sc
MAP calibrated date: 8016 BP
Te17
MAP calibrated date: 7975 BP
Ed2
MAP calibrated date: 7435 BP
Te23
MAP calibrated date: 7415 BP
Radiocarbon Age
Years cal BP

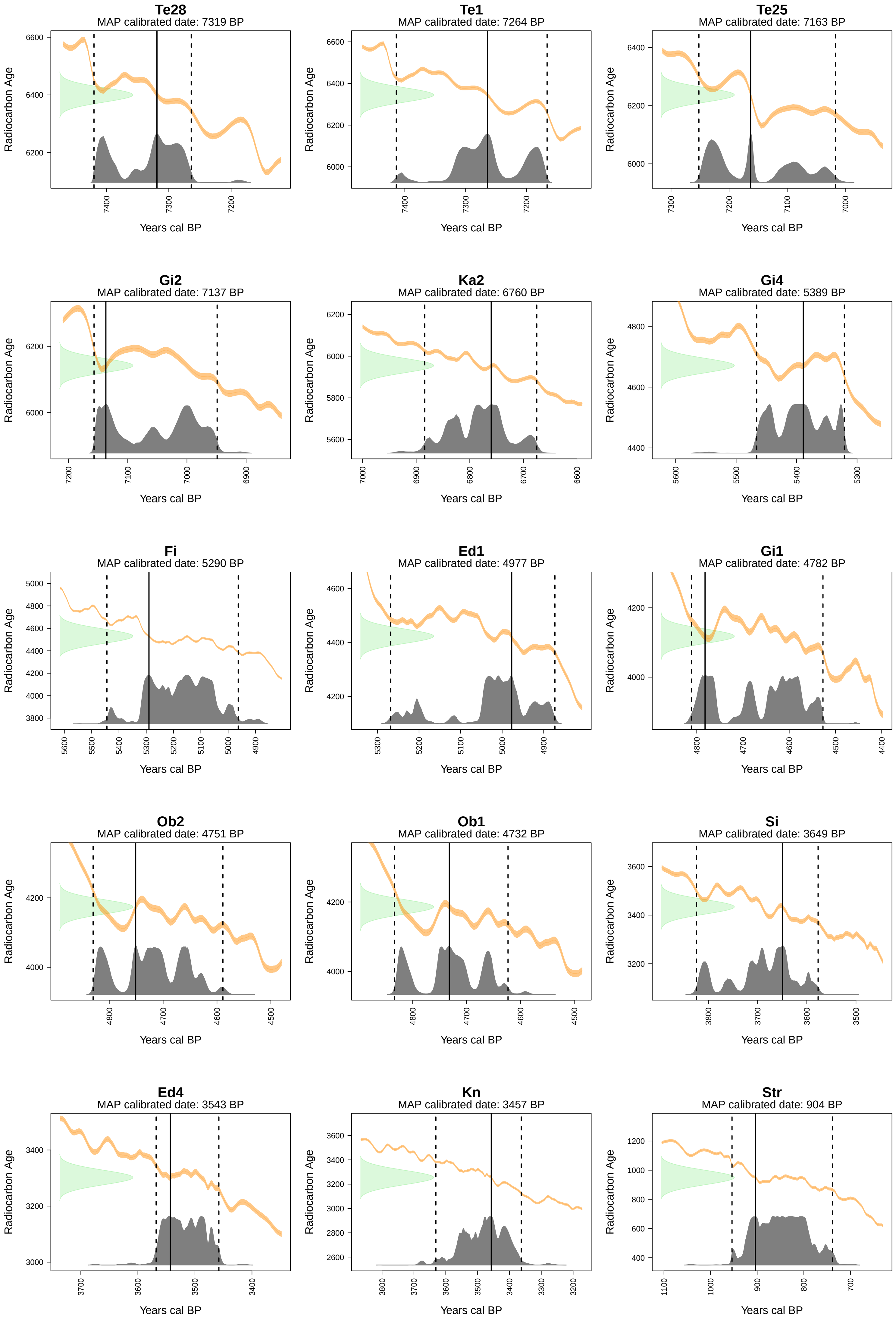
Te28
MAP calibrated date: 7319 BP
Te1
MAP calibrated date: 7264 BP
Te25
MAP calibrated date: 7163 BP
Gi2
MAP calibrated date: 7137 BP
Ka2
MAP calibrated date: 6760 BP
Gi4
MAP calibrated date: 5389 BP
Fi
MAP calibrated date: 5290 BP
Ed1
MAP calibrated date: 4977 BP
Gi1
MAP calibrated date: 4782 BP
Ob2
MAP calibrated date: 4751 BP
Ob1
MAP calibrated date: 4732 BP
Si
MAP calibrated date: 3649 BP
Ed4
MAP calibrated date: 3543 BP
Kn
MAP calibrated date: 3457 BP
Str
MAP calibrated date: 904 BP
Radiocarbon Age
Years cal BP

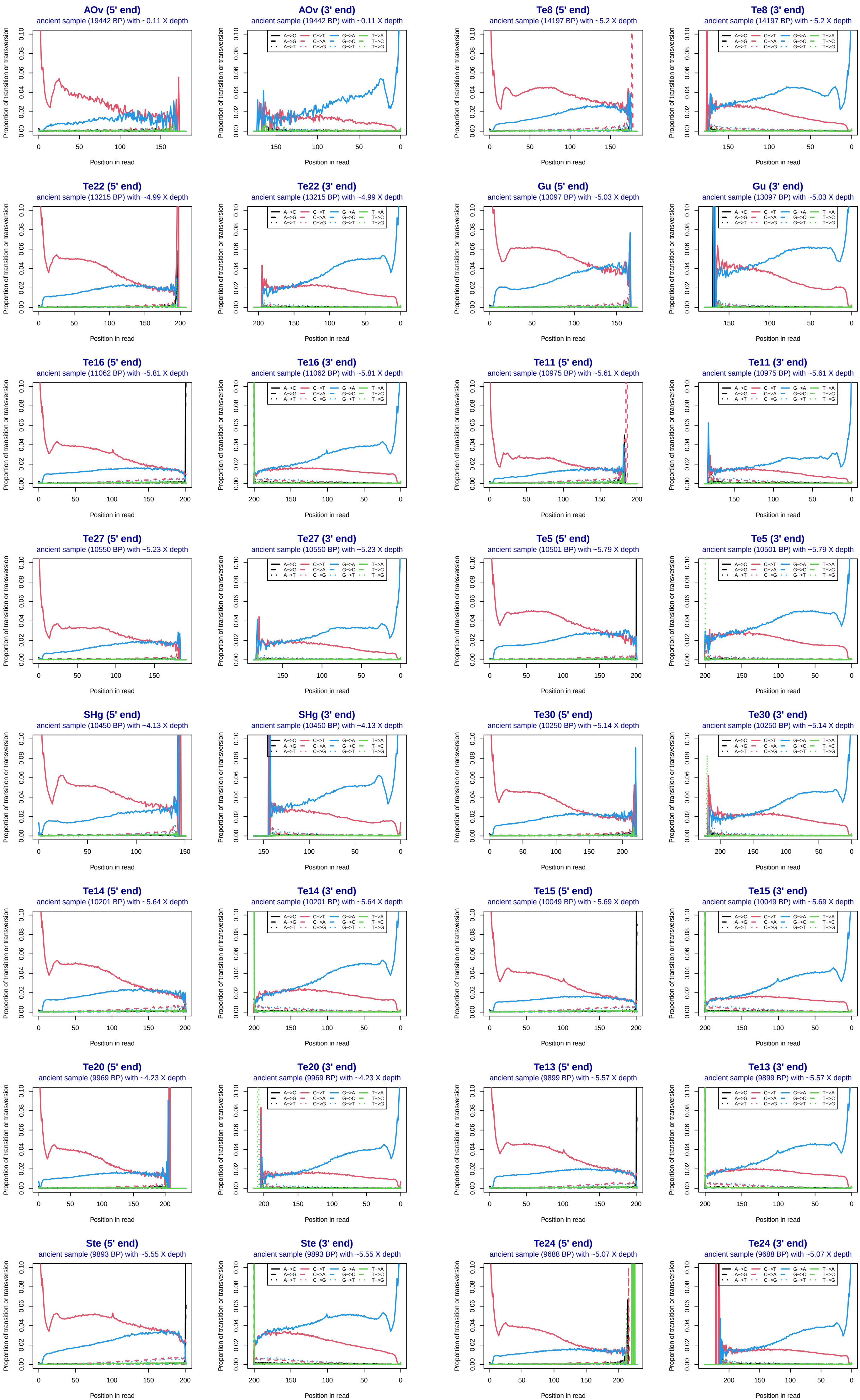
AOv (5' end)
ancient sample (19442 BP) with ~0.11 X depth
AOv (3' end)
ancient sample (19442 BP) with ~0.11 X depth
Te8 (5' end)
ancient sample (14197 BP) with ~5.2 X depth
Te8 (3' end)
ancient sample (14197 BP) with ~5.2 X depth
Te22 (5' end)
ancient sample (13215 BP) with ~4.99 X depth
Te22 (3' end)
ancient sample (13215 BP) with ~4.99 X depth
Gu (5' end)
ancient sample (13097 BP) with ~5.03 X depth
Gu (3' end)
ancient sample (13097 BP) with ~5.03 X depth
Te16 (5' end)
ancient sample (11062 BP) with ~5.81 X depth
Te16 (3' end)
ancient sample (11062 BP) with ~5.81 X depth
Te11 (5' end)
ancient sample (10975 BP) with ~5.61 X depth
Te11 (3' end)
ancient sample (10975 BP) with ~5.61 X depth
Te27 (5' end)
ancient sample (10550 BP) with ~5.23 X depth
Te27 (3' end)
ancient sample (10550 BP) with ~5.23 X depth
Te5 (5' end)
ancient sample (10501 BP) with ~5.79 X depth
Te5 (3' end)
ancient sample (10501 BP) with ~5.79 X depth
SHg (5' end)
ancient sample (10450 BP) with ~4.13 X depth
SHg (3' end)
ancient sample (10450 BP) with ~4.13 X depth
Te30 (5' end)
ancient sample (10250 BP) with ~5.14 X depth
Te30 (3' end)
ancient sample (10250 BP) with ~5.14 X depth
Te14 (5' end)
ancient sample (10201 BP) with ~5.64 X depth
Te14 (3' end)
ancient sample (10201 BP) with ~5.64 X depth
Te15 (5' end)
ancient sample (10049 BP) with ~5.69 X depth
Te15 (3' end)
ancient sample (10049 BP) with ~5.69 X depth
Te20 (5' end)
ancient sample (9969 BP) with ~4.23 X depth
Te20 (3' end)
ancient sample (9969 BP) with ~4.23 X depth
Te13 (5' end)
ancient sample (9899 BP) with ~5.57 X depth
Te13 (3' end)
ancient sample (9899 BP) with ~5.57 X depth
Ste (5' end)
ancient sample (9893 BP) with ~5.55 X depth
Ste (3' end)
ancient sample (9893 BP) with ~5.55 X depth
Te24 (5' end)
ancient sample (9688 BP) with ~5.07 X depth
Te24 (3' end)
ancient sample (9688 BP) with ~5.07 X depth
Proportion of transition or transversion
Position in read
A->C
A->G
A->T
C->T
C->A
C->G
G->A
G->C
G->T
T->A
T->C
T->G

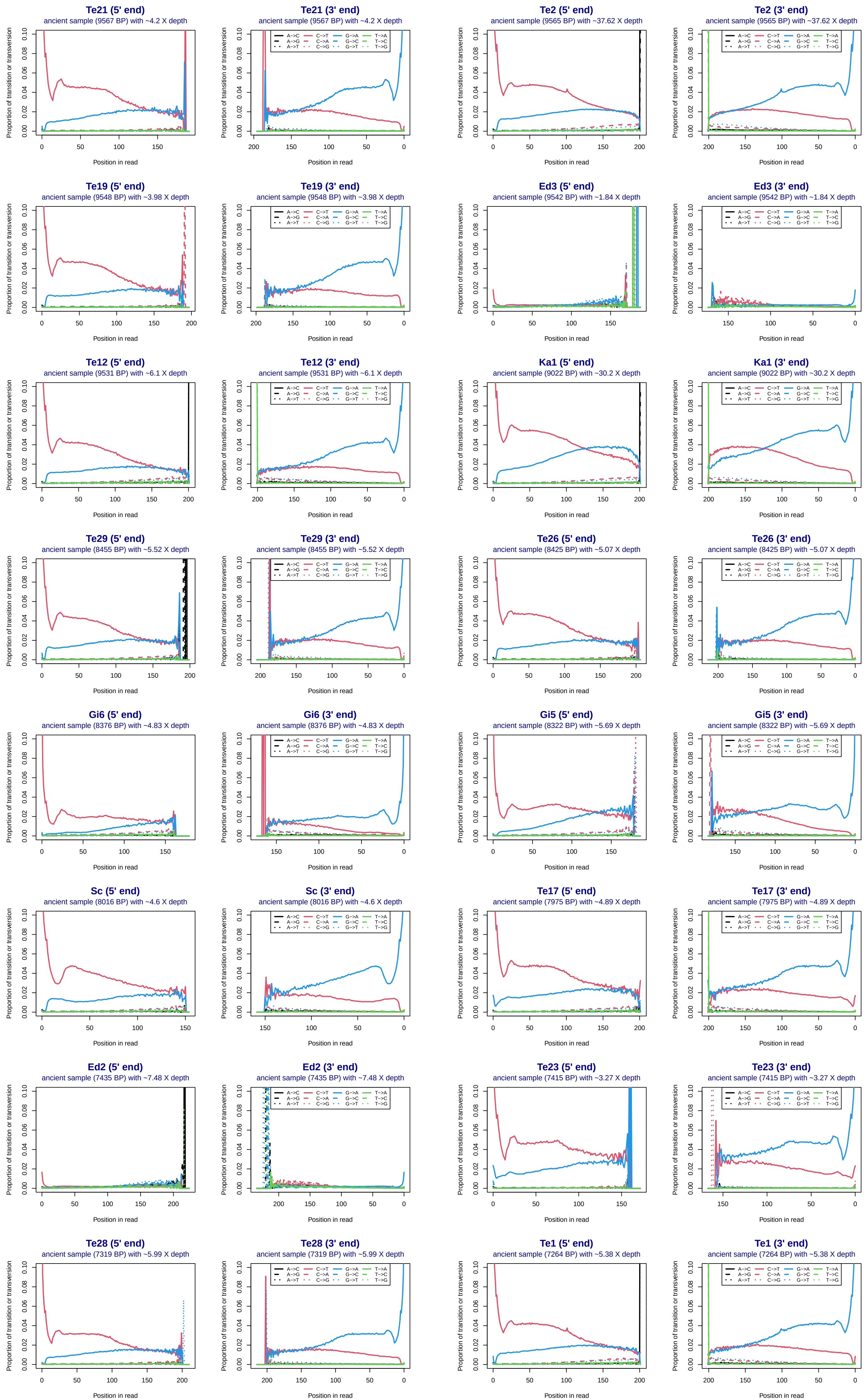
Te21 (5' end)
ancient sample (9567 BP) with ~4.2 X depth
Te21 (3' end)
ancient sample (9567 BP) with ~4.2 X depth
Te2 (5' end)
ancient sample (9565 BP) with ~37.62 X depth
Te2 (3' end)
ancient sample (9565 BP) with ~37.62 X depth
Te19 (5' end)
ancient sample (9548 BP) with ~3.98 X depth
Te19 (3' end)
ancient sample (9548 BP) with ~3.98 X depth
Ed3 (5' end)
ancient sample (9542 BP) with ~1.84 X depth
Ed3 (3' end)
ancient sample (9542 BP) with ~1.84 X depth
Te12 (5' end)
ancient sample (9531 BP) with ~6.1 X depth
Te12 (3' end)
ancient sample (9531 BP) with ~6.1 X depth
Ka1 (5' end)
ancient sample (9022 BP) with ~30.2 X depth
Ka1 (3' end)
ancient sample (9022 BP) with ~30.2 X depth
Te29 (5' end)
ancient sample (8455 BP) with ~5.52 X depth
Te29 (3' end)
ancient sample (8455 BP) with ~5.52 X depth
Te26 (5' end)
ancient sample (8425 BP) with ~5.07 X depth
Te26 (3' end)
ancient sample (8425 BP) with ~5.07 X depth
Gi6 (5' end)
ancient sample (8376 BP) with ~4.83 X depth
Gi6 (3' end)
ancient sample (8376 BP) with ~4.83 X depth
Gi5 (5' end)
ancient sample (8322 BP) with ~5.69 X depth
Gi5 (3' end)
ancient sample (8322 BP) with ~5.69 X depth
Sc (5' end)
ancient sample (8016 BP) with ~4.6 X depth
Sc (3' end)
ancient sample (8016 BP) with ~4.6 X depth
Te17 (5' end)
ancient sample (7975 BP) with ~4.89 X depth
Te17 (3' end)
ancient sample (7975 BP) with ~4.89 X depth
Ed2 (5' end)
ancient sample (7435 BP) with ~7.48 X depth
Ed2 (3' end)
ancient sample (7435 BP) with ~7.48 X depth
Te23 (5' end)
ancient sample (7415 BP) with ~3.27 X depth
Te23 (3' end)
ancient sample (7415 BP) with ~3.27 X depth
Te28 (5' end)
ancient sample (7319 BP) with ~5.99 X depth
Te28 (3' end)
ancient sample (7319 BP) with ~5.99 X depth
Te1 (5' end)
ancient sample (7264 BP) with ~5.38 X depth
Te1 (3' end)
ancient sample (7264 BP) with ~5.38 X depth
A->C C->T G->A T->A
A->G C->A G->C T->C
A->T C->G G->T T->G
Proportion of transition or transversion
Position in read

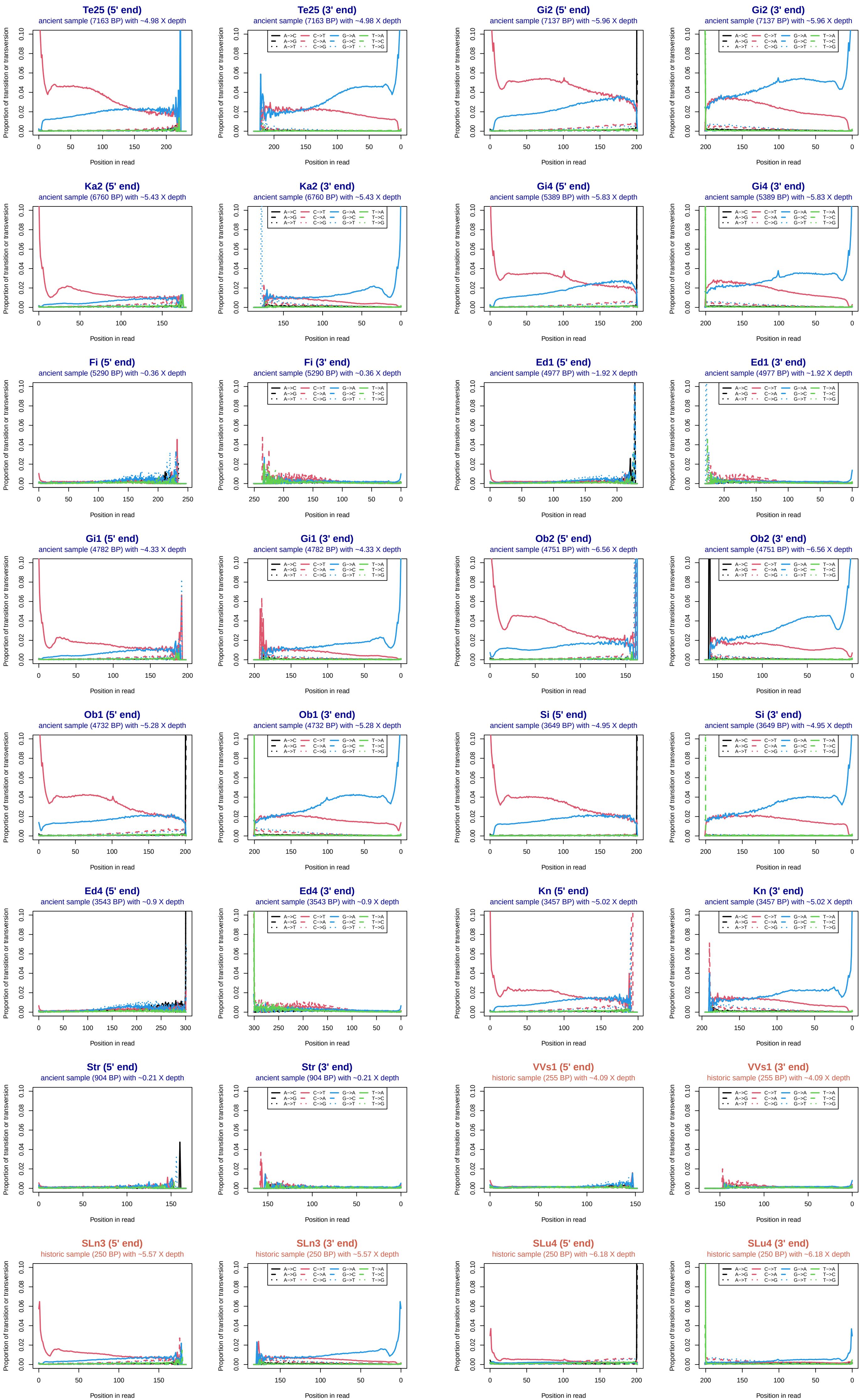
Te25 (5' end)
ancient sample (7163 BP) with ~4.98 X depth
Te25 (3' end)
ancient sample (7163 BP) with ~4.98 X depth
Gi2 (5' end)
ancient sample (7137 BP) with ~5.96 X depth
Gi2 (3' end)
ancient sample (7137 BP) with ~5.96 X depth
Ka2 (5' end)
ancient sample (6760 BP) with ~5.43 X depth
Ka2 (3' end)
ancient sample (6760 BP) with ~5.43 X depth
Gi4 (5' end)
ancient sample (5389 BP) with ~5.83 X depth
Gi4 (3' end)
ancient sample (5389 BP) with ~5.83 X depth
Fi (5' end)
ancient sample (5290 BP) with ~0.36 X depth
Fi (3' end)
ancient sample (5290 BP) with ~0.36 X depth
Ed1 (5' end)
ancient sample (4977 BP) with ~1.92 X depth
Ed1 (3' end)
ancient sample (4977 BP) with ~1.92 X depth
Gi1 (5' end)
ancient sample (4782 BP) with ~4.33 X depth
Gi1 (3' end)
ancient sample (4782 BP) with ~4.33 X depth
Ob2 (5' end)
ancient sample (4751 BP) with ~6.56 X depth
Ob2 (3' end)
ancient sample (4751 BP) with ~6.56 X depth
Ob1 (5' end)
ancient sample (4732 BP) with ~5.28 X depth
Ob1 (3' end)
ancient sample (4732 BP) with ~5.28 X depth
Si (5' end)
ancient sample (3649 BP) with ~4.95 X depth
Si (3' end)
ancient sample (3649 BP) with ~4.95 X depth
Ed4 (5' end)
ancient sample (3543 BP) with ~0.9 X depth
Ed4 (3' end)
ancient sample (3543 BP) with ~0.9 X depth
Kn (5' end)
ancient sample (3457 BP) with ~5.02 X depth
Kn (3' end)
ancient sample (3457 BP) with ~5.02 X depth
Str (5' end)
ancient sample (904 BP) with ~0.21 X depth
Str (3' end)
ancient sample (904 BP) with ~0.21 X depth
VVs1 (5' end)
historic sample (255 BP) with ~4.09 X depth
VVs1 (3' end)
historic sample (255 BP) with ~4.09 X depth
SLn3 (5' end)
historic sample (250 BP) with ~5.57 X depth
SLn3 (3' end)
historic sample (250 BP) with ~5.57 X depth
SLu4 (5' end)
historic sample (250 BP) with ~6.18 X depth
SLu4 (3' end)
historic sample (250 BP) with ~6.18 X depth
Proportion of transition or transversion
Position in read
A->C C->T G->A T->A
A->G C->A G->C T->C
A->T C->G G->T T->G

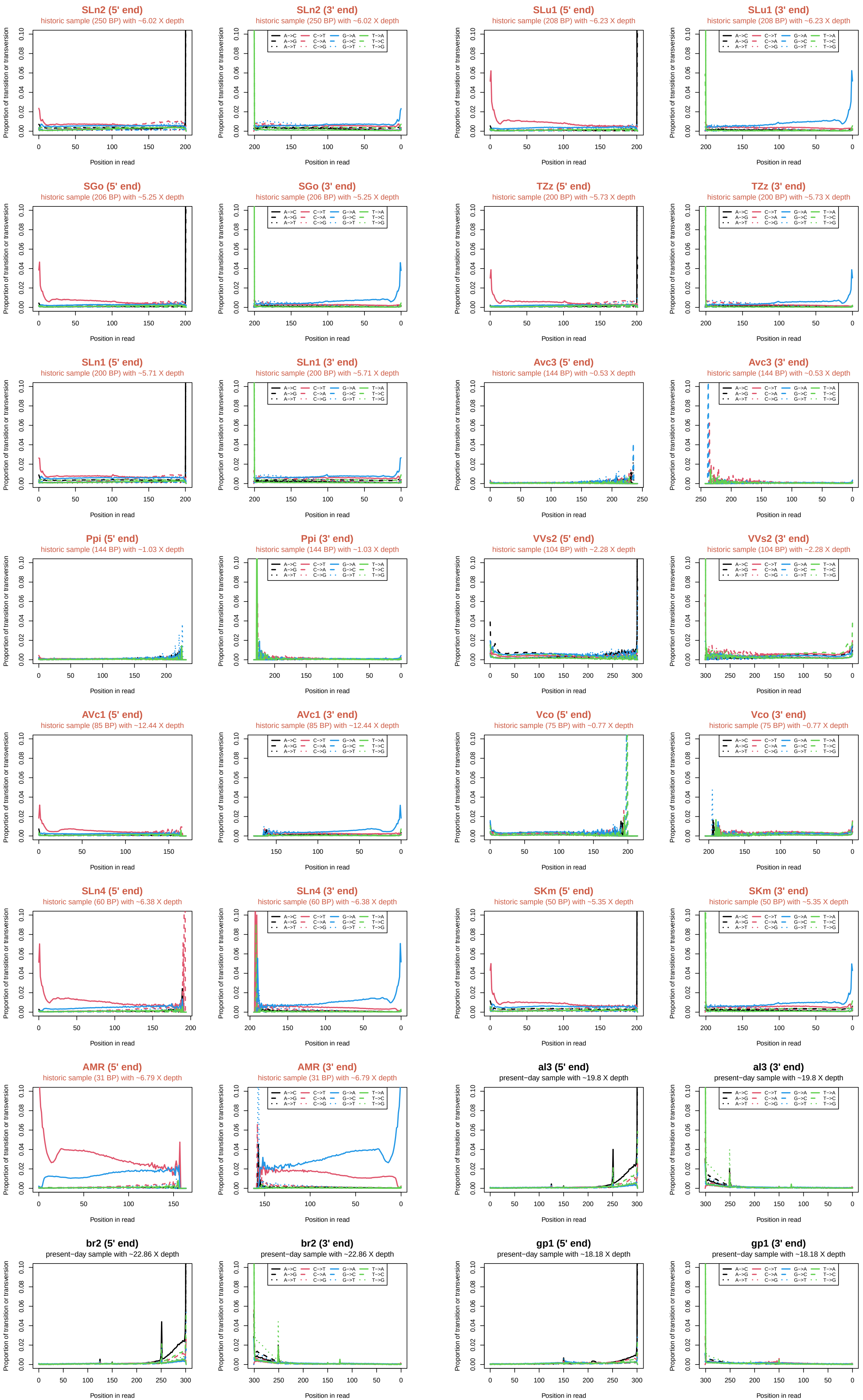
SLn2 (5' end)
historic sample (250 BP) with ~6.02 X depth
SLn2 (3' end)
historic sample (250 BP) with ~6.02 X depth
SLu1 (5' end)
historic sample (208 BP) with ~6.23 X depth
SLu1 (3' end)
historic sample (208 BP) with ~6.23 X depth
SGo (5' end)
historic sample (206 BP) with ~5.25 X depth
SGo (3' end)
historic sample (206 BP) with ~5.25 X depth
TZz (5' end)
historic sample (200 BP) with ~5.73 X depth
TZz (3' end)
historic sample (200 BP) with ~5.73 X depth
SLn1 (5' end)
historic sample (200 BP) with ~5.71 X depth
SLn1 (3' end)
historic sample (200 BP) with ~5.71 X depth
Avc3 (5' end)
historic sample (144 BP) with ~0.53 X depth
Avc3 (3' end)
historic sample (144 BP) with ~0.53 X depth
Ppi (5' end)
historic sample (144 BP) with ~1.03 X depth
Ppi (3' end)
historic sample (144 BP) with ~1.03 X depth
VVs2 (5' end)
historic sample (104 BP) with ~2.28 X depth
VVs2 (3' end)
historic sample (104 BP) with ~2.28 X depth
AVc1 (5' end)
historic sample (85 BP) with ~12.44 X depth
AVc1 (3' end)
historic sample (85 BP) with ~12.44 X depth
Vco (5' end)
historic sample (75 BP) with ~0.77 X depth
Vco (3' end)
historic sample (75 BP) with ~0.77 X depth
SLn4 (5' end)
historic sample (60 BP) with ~6.38 X depth
SLn4 (3' end)
historic sample (60 BP) with ~6.38 X depth
SKm (5' end)
historic sample (50 BP) with ~5.35 X depth
SKm (3' end)
historic sample (50 BP) with ~5.35 X depth
AMR (5' end)
historic sample (31 BP) with ~6.79 X depth
AMR (3' end)
historic sample (31 BP) with ~6.79 X depth
al3 (5' end)
present−day sample with ~19.8 X depth
al3 (3' end)
present−day sample with ~19.8 X depth
br2 (5' end)
present−day sample with ~22.86 X depth
br2 (3' end)
present−day sample with ~22.86 X depth
gp1 (5' end)
present−day sample with ~18.18 X depth
gp1 (3' end)
present−day sample with ~18.18 X depth
Proportion of transition or transversion
Position in read
A−>C A−>G A−>T C−>T C−>A C−>G G−>A G−>C G−>T T−>A T−>C T−>G

Figure S2: Post-mortem damage patterns. "Smile"-plot of relative transition and transversion proportions estimated using the ATLAS task refDiffTable for all sites passing site-level filters on scaffold 1. The forward (5' end) and the reverse (3' end) reads for each individual are plotted side by side. Present-day individuals are omitted, except for 6 individuals as examples.

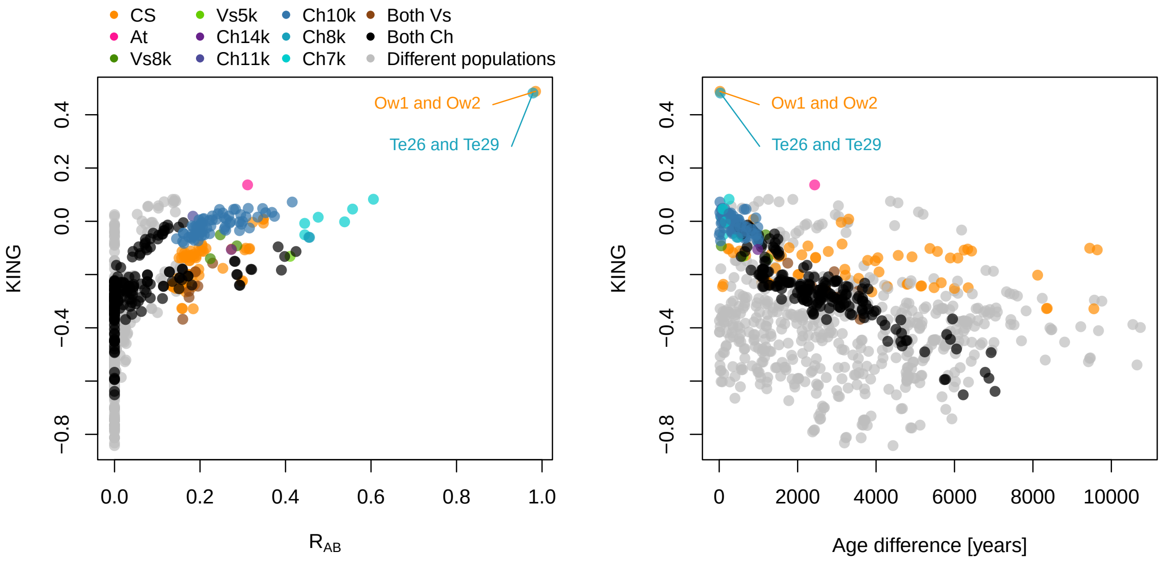


Figure S3: Pairwise relatedness. (A) Relatedness was estimated using both pairwise relatedness (RAB) and the KING kinship coefficient in ngsRelate v2.3.0 (Korneliussen and Moltke, 2015). The scatterplot shows pairwise relatedness estimates for 42 ancient Alpine ibex samples. Using the midpoint between the theoretical estimates of pairwise relatedness as in Seersholm et al., 2024, pairs are classified as first-degree relatives with a RAB between 0.75 and 0.25. An RAB $>$ 0.75 indicates identical individuals. 1 (out of 66) pairs of the population Ch10k are classified as first-degree relatives as well as 2 (out of 3) pairs of population Ch8k, 6 (out of 6) pairs of population Ch7k and the pair of Vs5k individuals. The two pairs of individuals classified as identical are specifically noted in the plot as Ow1 and Ow2 (population CS) and Te26 and Te29 (population Ch8k). Using the inference criteria (Manichaikul et al., 2010) of the kinship coefficient (KING) (Waples, Albrechtsen, and Moltke, 2019), no sample pairs are categorised as first-degree relatives and the two pairs noted in the plot are again categorised as identical. (B) There is no effect of sampling age on the relatedness of individuals.

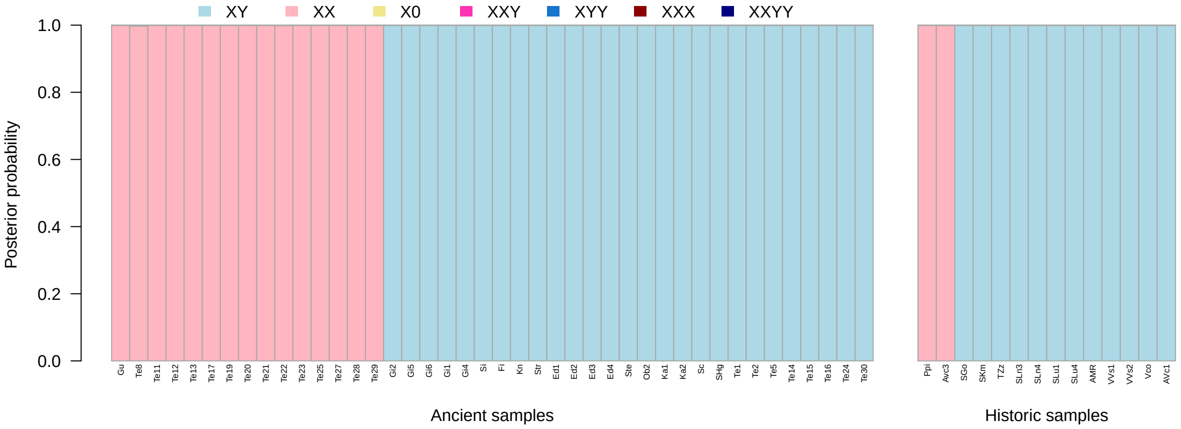


Figure S4: Sex identification. Posterior probability of individual sex karyotypes inferred by BeXY (Caduff et al., 2024). All samples were classified with high confidence as XX (n = 26) or XY (n = 39).

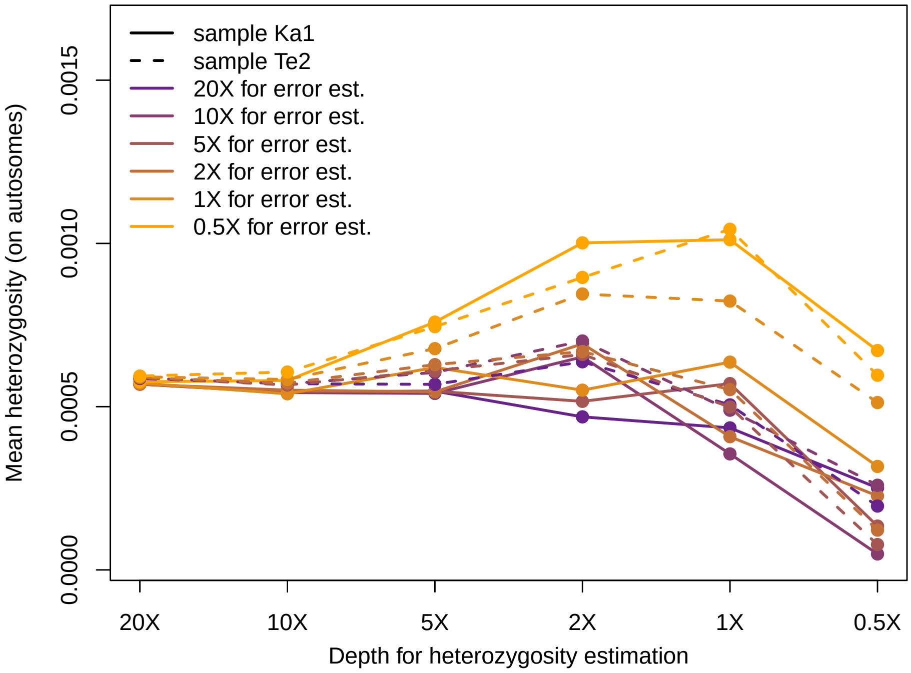


Figure S5: Downsampling experiment using two ancient Alpine ibex samples sequenced to high depth (Ka1 and Te2). The displayed statistic is the heterozygosity estimated under the HKY85 genotype substitution model (Hasegawa, Kishino, and Yano, 1985). Downsampling on error estimation level is displayed as colour and downsampling at heterozygosity estimation level is displayed on the X-axis.

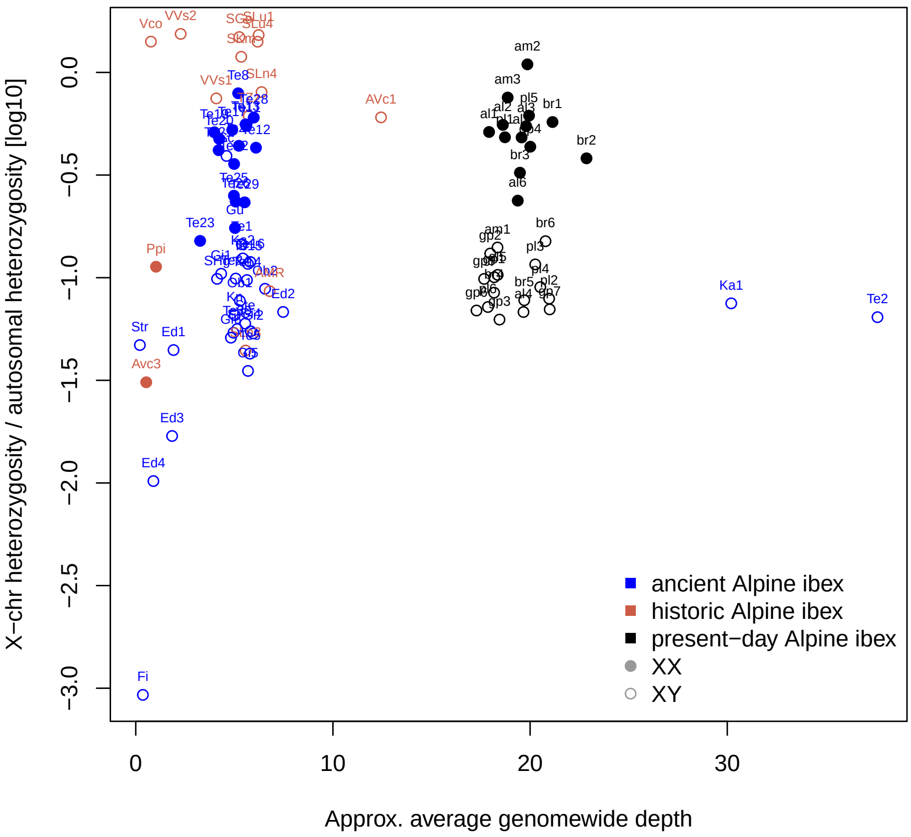


Figure S6: Relative heterozygosity on the X chromosome compared to the autosome.

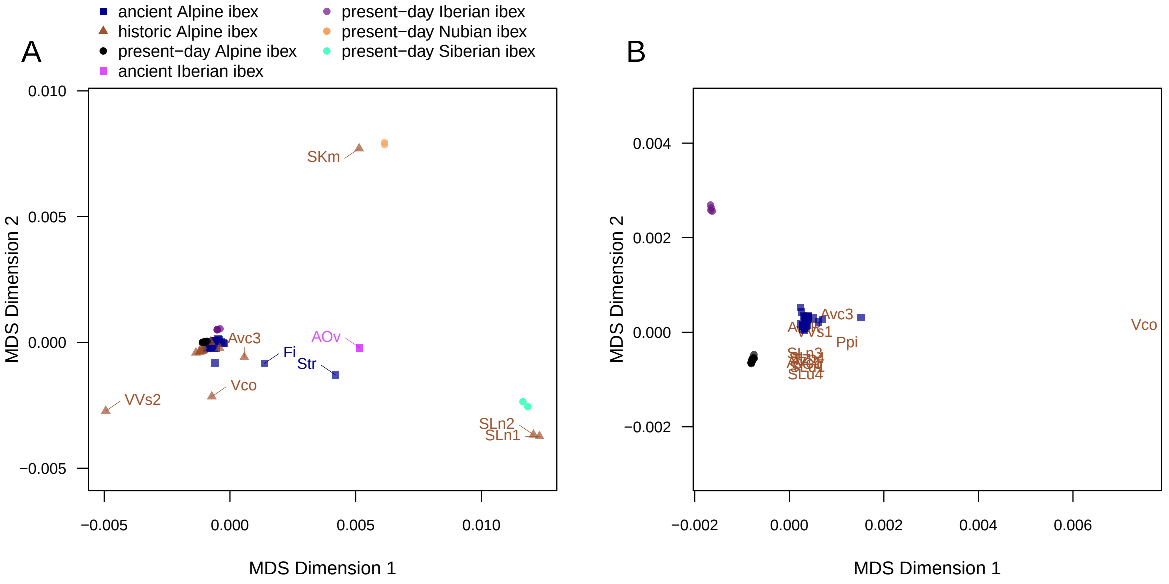


Figure S7: (A) Multi-dimensional scaling (MDS) of the genetic distances between all pairs of ibex samples. Outlier historic and ancient samples are indicated. (B) MDS of genetic distances of individuals clustering with present-day Alpine ibex samples and excluding samples clustering with different ibex species.

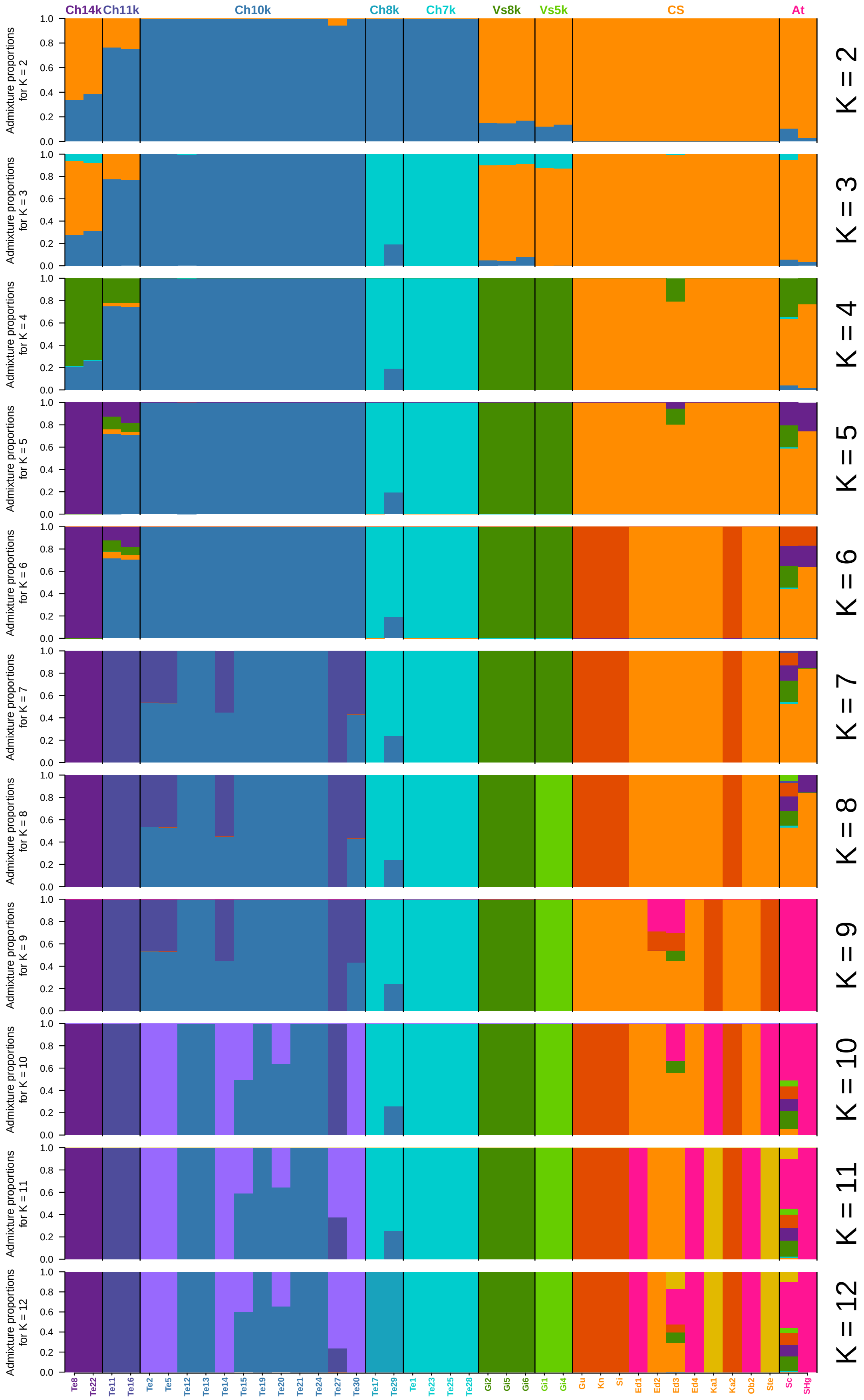


Figure S8: Unsupervised genetic clustering of ancient Alpine ibex individuals with ngsAdmix for 2-12 clusters (K). Results from the iteration with the highest log likelihood (out of 20 repetitions) are shown for each K-value.

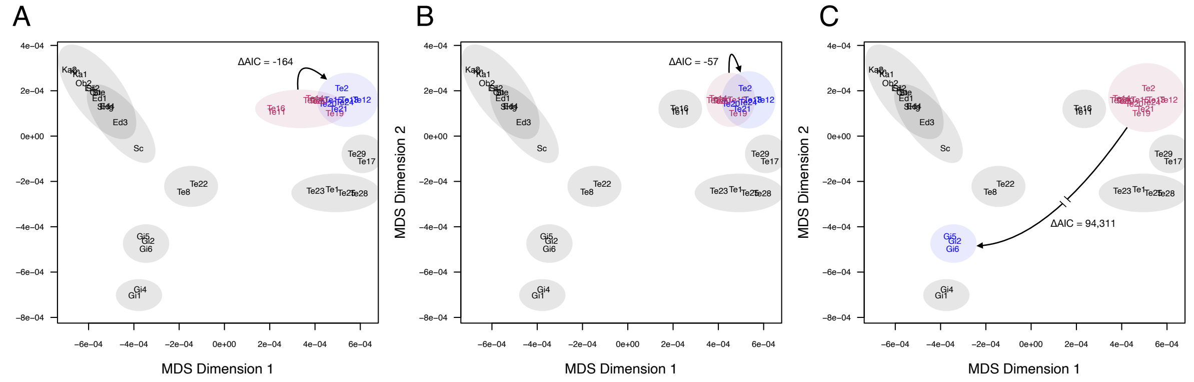

Figure S9: Continuity tests performed in `fastsimcoal2` (Excoffier et al., 2021) to test whether model holds up to expected result in test scenarios focusing on the sample-rich cluster Ch10k. (A) Testing for continuity when both populations Ch11k and Ch10k are combined and then split into two with the younger 7 samples in the blue population and the older 7 samples in the red population. The model for a continuous population outperforms the population turnover model (negative ΔAIC). (B) Similar test for continuity when population Ch10k is split into two with the younger 6 samples in the blue population and the older 6 samples in the red population. (C) Test for continuity between two distant populations Vs8k (in blue) and Ch10k (in red). The model for a population turnover outperforms the model for a continuous population with a very big margin (strongly positive ΔAIC).

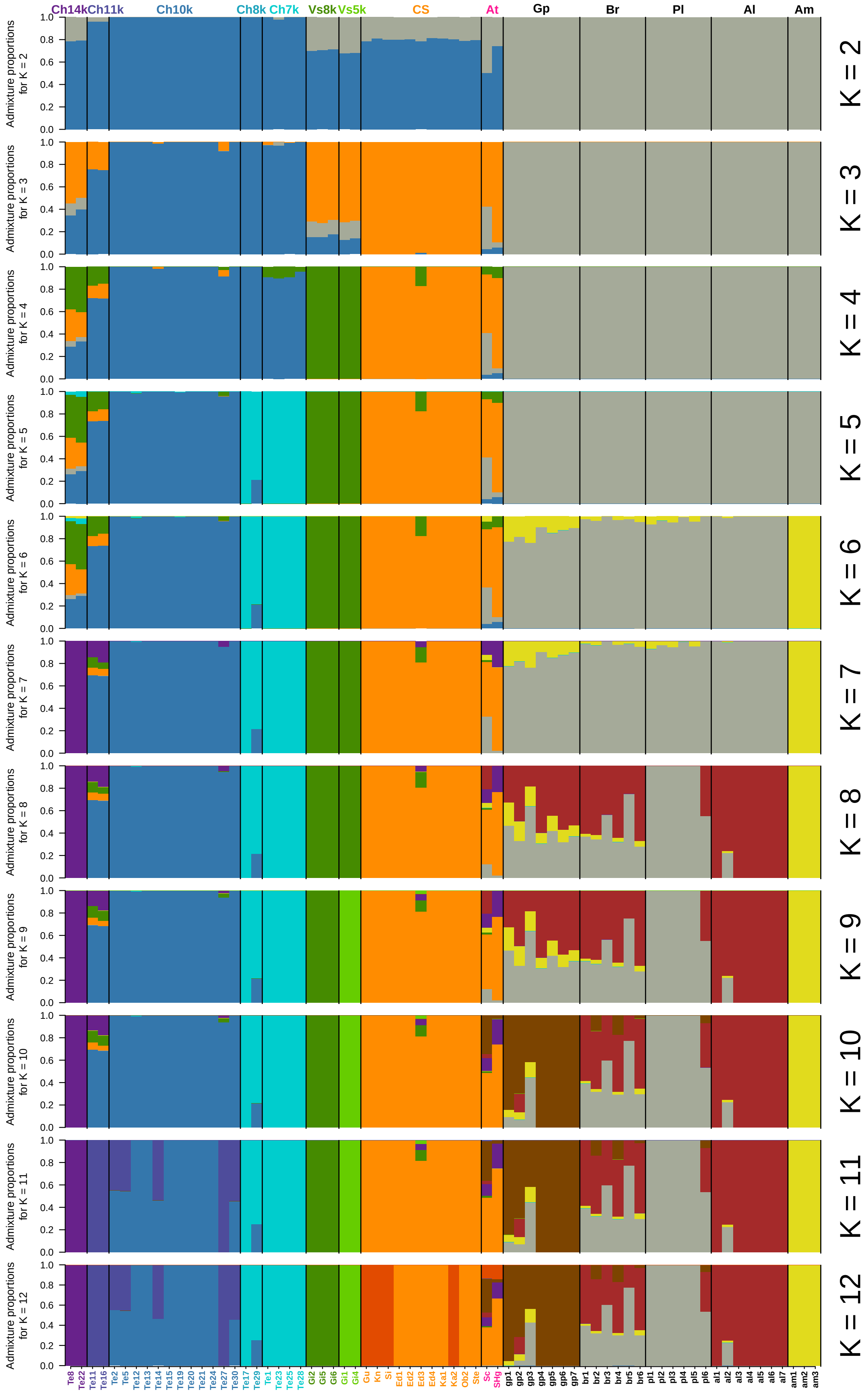


Figure S10: Unsupervised genetic clustering of ancient and present-day Alpine ibex individuals with ngsAdmix for 2-12 clusters (K). Results from the iteration with the highest log likelihood (out of 20 repetitions) are shown for each K-value.
